# High-Definition MEG Source Estimation using the Reciprocal Boundary Element Fast Multipole Method

**DOI:** 10.1101/2025.03.21.644601

**Authors:** Guillermo Nuñez Ponasso, Derek A. Drumm, Abbie Wang, Gregory M. Noetscher, Matti Hämäläinen, Thomas R. Knösche, Burkhard Maess, Jens Haueisen, Sergey N. Makaroff, Tommi Raij

## Abstract

*Magnetoencephalographic* (MEG) source estimation relies on the computation of the gain (lead-field) matrix, which embodies the linear relationship between the amplitudes of the sources and the recorded signals. However, with a realistic forward model, the calculation of the gain matrix in a “direct” fashion is a computationally expensive task because the number of dipolar sources in standard MEG pipelines is often limited to ∼10,000. We propose a fast approach based on the reciprocal relationship between MEG and transcranial magnetic stimulation (TMS). This approach couples naturally with the charge-based boundary element fast multipole method (BEM-FMM), which allows us to efficiently generate gain matrices for high-resolution multi-layer non-nested meshes involving source spaces of up to a ∼1 million dipoles. We evaluate our approach by performing MEG source reconstruction against simulated data (at varying noise levels) obtained from the direct computation of MEG readings from 2000 different dipole positions over the cortical surface of 5 healthy subjects. Additionally, we test our methods with real MEG data from evoked somatosensory fields by right-hand median nerve stimulation in these same 5 subjects. We compare our experimental source reconstruction results against the standard MNE-Python source reconstruction pipeline.

## 1 Introduction

Source estimation (Knösche and Haueisen 2022) aims to estimate neural activity sites and time courses from non-invasive measurements such as *electroencephalography* (EEG) and *magnetoencephalography* (MEG) (Hämäläinen et al. 1993). This technique can be used clinically to diagnose pathologies, for example, by localizing sources of interictal epileptic activity (Plummer et al. 2008), or studying ADHD (Jonkman et al. 2004); it also has basic-research applications in mapping higher cognitive function (Siebenhühner et al. 2016; Gross 2019), in psychological and psychiatric research (Foti et al. 2011; Hämäläinen et al. 2015; Uhlhaas et al. 2017), and as a component in brain-computer interfaces (Noirhomme et al. 2008).

The methodology of MEG/EEG source estimation can be split into two main components: the forward model and the inverse model. The forward model depends on subject anatomy, sensor placement and specifications, and assumptions on conductivity values. In this step, sensor outputs are calculated for each individual putative source, each modeled as a current dipole (Hämäläinen et al. 1993), by solving the quasistatic Maxwell equations (Nuñez Ponasso 2024). An important point is that the only sources of indeterminacy in the forward stage are due to MRI segmentation errors (Lanfer et al. 2012), uncertainty of conductivity values (Homma et al. 1995; Haueisen et al. 1999; Vorwerk et al. 2024), and co-registration errors (Chella et al. 2019); any other errors are numerical.

On the other hand, the inverse problem is ill posed. Even though an infinite number of possible solutions exist (Helmholtz 1853), useful and accurate estimates of the source locations can be obtained by introducing biophysical constraints. Examples of such assumptions are: 1) limiting the source model to the interface between gray and white matter; 2) assuming that equivalent current dipoles are oriented perpendicular to the gray matter surface (Nunez 1981); or 3) assumptions on the focality or smoothness (spread) of the solution based on the experimental paradigm (Schmidt et al. 1999).

While such assumptions are reasonable, computational constraints impose artificial assumptions like limiting the number of sources or reducing the number of compartments in the head model. The MNE software (Gramfort et al. 2013) is one of the most widely used freely available source reconstruction packages. Source estimation on MNE is based on the potential-based BEM formulation of the MEG forward model (Geselowitz 1967; Geselowitz 1970; Mosher et al. 1999; Nuñez Ponasso 2024); because of this, the forward system matrix is dense and the method is often limited to 3 compartments: the skin, skull, and pial surface. Furthermore, it is often limited to ∼ 20 000 possible sources (Stenroos et al. 2014) even though using denser source spaces is possible with additional computational cost. These limitations are present in most source-reconstruction software like MNE (Gramfort et al. 2013; Gramfort et al. 2014), FieldTrip (Oostenveld et al. 2010), Brainstorm (Tadel et al. 2011), or SPM (SPM 2025). As a general principle, we should remove such limitations and base our inverse model on forward solutions that model the underlying bioelectromagnetics as accurately as possible (Dale and Sereno 1993).

The main ingredient tying the forward model and the inverse model is the so-called *leadfield matrix*. This is a matrix of size (*M× N*), where M is the number of sensors and N is the number of sources, and *N >> M* . Each column of the leadfield matrix. The major computational limitation to the number of sources is that, in the classical direct approach, the columns of the matrix must be computed one at a time.

The *charge-based boundary element fast multipole method* (or BEM-FMM) (Greengard et al. 2009) is a novel technique for forward computations in bioelectromagnetism noted for its speed, accuracy, and ability to deal with high-resolution meshes with non-nested topology (Makarov et al. 2018; Makarov et al. 2021). In Makarov et al. 2020, a BEM-FMM toolkit to solve *transcranial magnetic stimulation* (TMS) forward problems was proposed. Due to *Lorentz’s reciprocity formula* (Heller and Hulsteyn 1992), the forward MEG problem can be solved by means of the forward TMS problem rather than computing forward solutions for equivalent current dipoles. Hence, we solve a TMS forward solution where the sensor coils act as TMS coils. We use Lorentz’s formula to “fill” the leadfield matrix by its rows instead of by its columns, which is the *reciprocal approach*. The advantage is clear: while the number of sources (columns) can be in the millions, the number of sensors (rows) is no larger than a few hundred in a typical MEG system. Furthermore, the coupling between BEM-FMM and the reciprocal MEG computation has two additional advantages: 1) it is sufficient to compute the charge densities induced by the sensors as TMS coils; 2) since in the TMS forward problem no singularities are present near the computational meshes, there is no need for *adaptive mesh refinement* (AMR) (Weise et al. 2022; Wartman et al. 2024) in order to calculate the TMS solutions. This is in contrast to the direct computation of high-resolution EEG or MEG, which often requires AMR to improve numerical stability (Nuñez Ponasso et al. 2024; Wartman et al. 2025), which adds to the computational cost.

We tested our proposed source reconstruction method by performing localization of *somatosensory evoked fields* elicited by right-hand median nerve stimulation and comparing our results with the output of the standard MNE pipeline using MNE-Python. To ensure that differences in the solutions are only due to the difference in forward method, we replicated one-to-one the inverse methods employed by MNE. Evoked somatosensory fields are known to be generated in the Brodmann area 3b (Purves et al. 2004; Brodmann 2005)in the primary somatosensory cortex situated in the posterior wall of the central sulcus (Nuwer 1998), so these fields provide a good testing ground. In addition, the localization of evoked somatosensory fields is also interesting for clinical applications (Lüders et al. 1986).

We also tested our inverse methods against simulated MEG data and, to avoid possible biases caused by using the same forward operator, we generated MEG signals using direct (non-reciprocal) BEM-FMM with *b*-refinement (Wartman et al. 2024). We compared distances from the peaks and centroids of the estimated regions of activation to the ground-truth source dipole location at various noise levels.

## 2 Materials and methods

### 2.1 Reciprocal construction of the MEG leadfield matrix

The Lorentz reciprocity theorem (Supplement A) establishes a relation between two formally independent fields operating at the same frequency *ω*,the field of a given dipole and the field of an arbitrary MEG sensor (magnetometer or gradiometer) operating as an induction (e.g. TMS) coil. In Supplement A it is shown that if both fields are assumed to be in phase, the reciprocity relation can be written in the following real form (without using complex phasors):

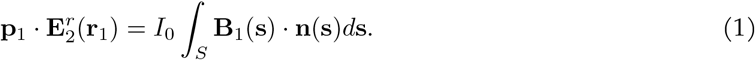

Here, indices with value 1 refer to the current dipole, and indices with value 2 refer to the MEG sensor coil. Our notations are as follows:

– **p**_1_ [A m] is the vector dipole moment of the primary current dipole source located at **r**_1_ with current strength **J**_1_(**r, r**) = **p**_1_*δ*(**r** − **r**_1_) cos *ωt*.
– *I*_0_ [A] is the amplitude of a harmonic coil current **I**(*t*) = *I*_0_ cos(*ωt*) flowing through the MEG sensor coil.
– **B**_1_ [T] is the spatial distribution of the magnetic flux density **B**_1_(**r**; *t*) = **B**_1_(**r**) cos(*ωt*) generated by the current dipole. The integral term is the magnetic flux through the surface *S* bound by the MEG coil.
– 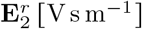 is the spatial distribution of the out-of-phase total electric field 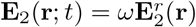 sin *ωt* generated by the MEG sensor operating as an induction coil, up to a multiplicative factor *ω* [s^−1^].

The frequency *ω* can be chosen arbitrarily. Equation 1 coincides with Equation (1) of (Nummenmaa et al. 2013) when the harmonic coil current is selected as stated above.

### 2.2 Multiple dipoles and one MEG sensor

It is shown in Plonsey 1972 that the quasi-static approximation to Maxwell’s equations can be used for EEG and MEG modeling. Equation 1 can then be generalized to multiple dipoles and multiple MEG sensors using linearity. We consider multiple dipoles **p**_*j*_, *j* = 1, …, *N* . We may assume that the dipole moments are perpendicular to the cortex (Hämäläinen et al. 1993), i.e. **p**_*j*_ = *p*_*j*_**n**_*j*_, where **n**_*j*_ is the unit normal vector to the cortical surface at **r**_*j*_. These dipoles generate the net static magnetic field **B**(**r**) = **B**_1_(**r**)+… +**B**_*N*_ (**r**) obtained by superposition. Applying Equation 1 to each dipole individually, and summing, yields:

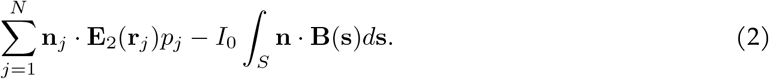

### 2.3 Multiple dipoles and multiple MEG sensors

We turn on only one *i*-th induction coil (one sensor) at a time while leaving all others turned off. We assume the same current amplitude *I*_0_ for every coil (sensor). For *M* sensors, this will give us *M* equations like Equation 2 by running over all sensors (coils) when *i* = 1, …, *M* . These equations can be written in compact matrix form (omitting indices 1 and 2 previously used):

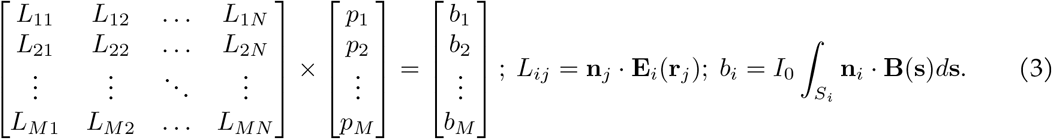

The matrix on the left-hand side of Equation 3 is denoted by 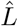.This is indeed the standard unique MEG leadfield matrix which relates unknown dipole strengths *p*_*j*_, *j* = 1, …, *N* to the measured sensor responses *b*_*i*_, *i* = 1, …, *M* — a discretized representation of the forward problem (Knösche and Haueisen 2022). The factor *I*_0_ in Equation 3 is superfluous; it can be chosen to be 1 A s^−1^. However, in contrast to the classical leadfield matrix definition (Knösche and Haueisen 2022), the present formula explicitly specifies elements *L*_*ij*_ of the leadfield matrix in terms of the reciprocal fields **E**_*i*_ as, for example, in Nummenmaa et al. 2013 and Nolte 2003.

### 2.4 Comparison of direct and reciprocal approaches to constructing the lead field matrix

A direct approach widely used today in the boundary element method (BEM) software packages such as MNE (Gramfort et al. 2013), Brainstorm (Tadel et al. 2011), and FieldTrip (Oostenveld et al. 2010) does not employ electric fields of the MEG coils. Instead, magnetic fields of every dipole, **B**_1_, **B**_2_, *· · ·*, **B**_*N*_, which are present on the right-hand side of Equation 3, are computed directly at all sensor locations for unit-strength dipoles at all putative source locations. In other words, 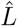 is filled *column-wise*, for every dipole separately. The number of such computations is equal to the number of dipoles *N* . We note that *N* ≫ *M* .

A currently less used alternative approach is a “reciprocal” BEM (Nummenmaa et al. 2013; Wendel et al. 2009; Laarne et al. 2000), where the lead field matrix is defined in terms of the reciprocal fields **E**_*i*_. In other words, 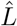 is filled *row-wise*. We excite a single sensor at a time, compute the corresponding induced field everywhere in the cortex, e.g. at all dipole positions **r**_1_, **r**_2_, …, **r**_*N*_, find the dot product of this field with the normal cortical vectors **m**_1_, **m**_2_, …, **m**_*N*_, and finally obtain the complete *i*-th row of 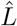. The number of such computations is equal to the number of sensors *M* .

### 2.5 Expansion of a cortical dipole density into global cortical basis functions

The reciprocal approach for a dense source space can be interpreted as an expansion of the unknown continuous dipole density *p*(**r**) into a set of global basis functions, defined over the entire cortical surface, similar to spatial Fourier harmonics. To within a normalization constant, every such basis function is simply an electric field 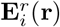 of an MEG sensor (gradiometer or magnetometer) operating as a TMS coil and computed at a cortical surface of interest in Equation 3. For the normal cortical dipole density, it is given by the projection of the electric field onto the normal cortical direction, 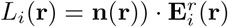 where **r** belongs to the cortical surface of interest. Thus, we have

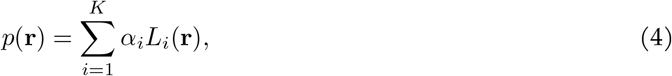

with unknown coefficients *α*_*i*_. The expansion in Equation 4 is performed separately for two sets of perpendicular gradiometers and one set of magnetometers. The total number of unknown coefficients is therefore the total number of sensors. Figure 1 illustrates the (normalized) basis functions *L*_*i*_(**r**) for two gradiometers and one magnetometer, respectively, computed for the first subject of a cohort described further in the text. The cortical surface chosen is the surface just outside the white matter resulting from a small shift of the white matter surface in the direction towards the grey matter surface.

**Figure 1.**
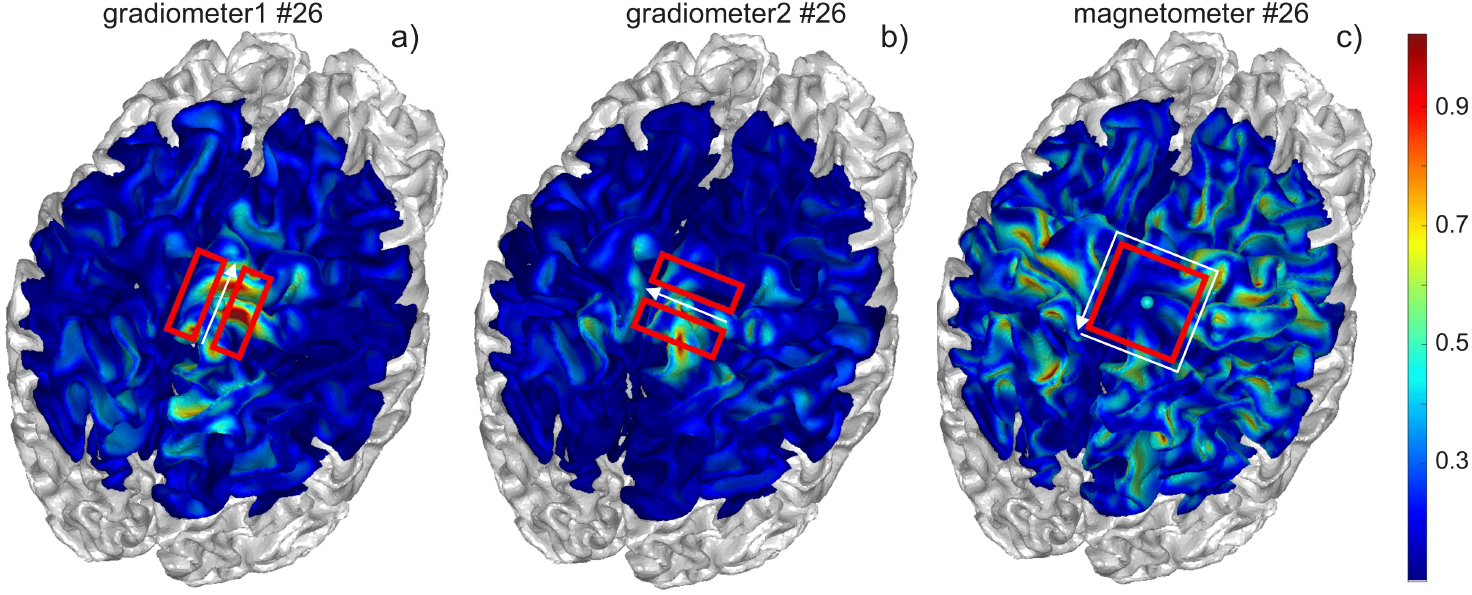
Normalized basis functions *L*_*i*_(**r**) for two perpendicular gradiometers and one magnetometer, respectively, computed for the first subject of a cohort described further in the text. The dominant current direction in the TMS mode is shown by the white arrow. The cortical surface chosen is the surface just outside white matter. The gradiometer basis functions are well localized over the cortical surface where the magnetometer basis functions are not.

### 2.6 Accelerated computation of normal component of E-field via charge densities

Since BEM-FMM computes surfaces charges, we can obtain the normal component of the total electric field without computing the secondary field (Makarov et al. 2020). Let *c* be the surface charge density and let be a point at the WM-GM interface, then we have:

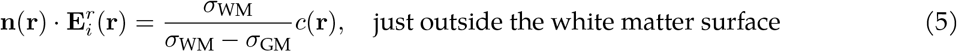

Similarly, if **r** is a point at the GM-CSF interface, then we have

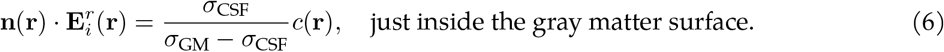

This is in contrast to the finite element method (FEM) or the surface potential formulation of the BEM, where the calculation of the secondary field is necessary. In the following, we will choose the cortical surface as the one just outside the white matter interface and then use Equation 5 directly.

### 2.7 Forward TMS computations using the FMM-accelerated charge-based boundary element method

To compute the forward TMS solution, we apply the charge-base boundary element fast multipole method (Greengard et al. 2009; Makarov et al. 2018). This method uses the charge-based formulation of the boundary element method, where the charge density *ρ* induced by an arbitrary incident field **E**^*i*^ is computed according to the following integral equation (Makarov et al. 2018; Nuñez Ponasso 2024):

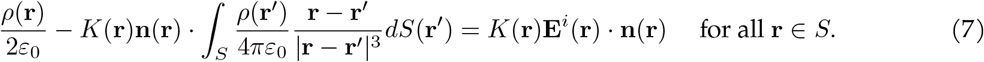

Here, we are assuming that the conductivity *σ* is isotropic, and that *S* is the surface of discontinuity of *σ*. The vector **n**(**r**) is the outward normal to the surface *S* at a point **r** of *S*. The constant *ε*_0_ ≈ 8.8541 *×* 10^−12^ [F m^−1^] is the permittivity of free space. The dimensionless scalar *K*(**r**) is the *conductivity contrast* at an interface point **r** of *S*, defined as

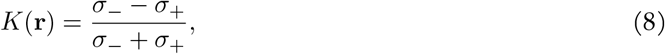

where *σ*_+_ (*σ*_−_) is the conductivity just outside (just inside) of the surface *S* at the point **r**.

In our case, the incident electric field is the one generated by a current loop of unit strength following the outline of the MEG sensors; this is equivalent to calculating a TMS solution using the sensors as stimulation coils. To model the incident field **E**^*i*^ generated by a TMS coil, we use a series of current elements (Makarov et al. 2020) of current *i*_*j*_(*t*) with orientation vector **s**_*j*_ and center **p**_*j*_, for *j* = 1, …, ℓ. These current elements can be distributed uniformly across the cross-section of the coil to model Litz wires, or non-uniformly for wires susceptible to the skin effect. The primary field 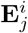 produced by the *j*-th current element is given by (Balanis 2012; Makarov et al. 2020),

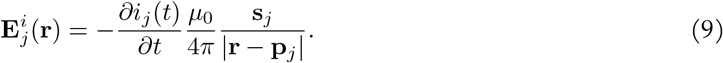

Where *µ*_0_ ≈1.2566 *×* 10^−6^ N A^−2^ is the permeability of free space. The primary field is calculated by aggregating the contribution of each current element:

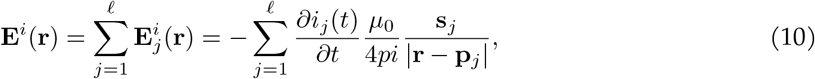

and this summation term can be accelerated using the fast multipole method (FMM) (Greengard et al. 2009).

Using the Galerkin method, the charge-based Equation 7 becomes discretized as:

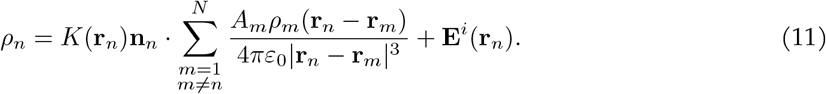

The solution of Equation 11 can be accelerated by using FMM to compute the summation term. Using the first-order approximation 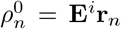,Equation 11 can be used to iteratively solve for *ρ*_*n*_ (Nuñez Ponasso 2024): At every step *k* — having computed the successive approximations 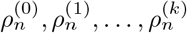 — we compute a *generalized minimimum residual* (GMRES) solution by calculating the parameters *α*_0_, *α*_1_, …, *α*_*k*_ minimizing the residual in the expression

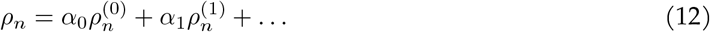

The chosen precision parameters for the BEM-FMM solution using GMRES are indicated in Table 1.

**Table 1:**
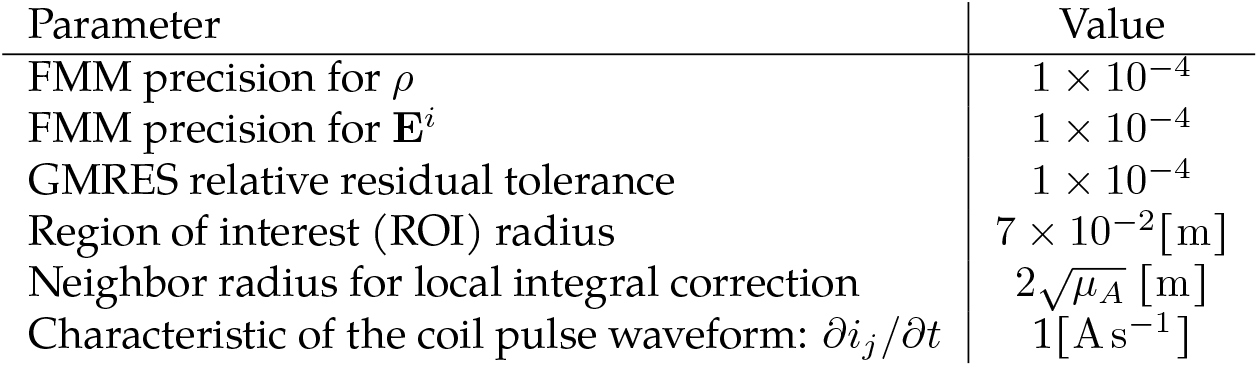
Parameters selected for the TMS forward computations. The symbol *µ*_*A*_ denotes the average triangle area in the mesh. The absence of a unit label in the “Value” column indicates a dimensionless parameter.

| Parameter | Value |
| --- | --- |
| FMM precision for $\rho$ | $1 \times 10^{-4}$ |
| FMM precision for $\mathbf{E}^i$ | $1 \times 10^{-4}$ |
| GMRES relative residual tolerance | $1 \times 10^{-4}$ |
| Region of interest (ROI) radius | $7 \times 10^{-2}[\text{m}]$ |
| Neighbor radius for local integral correction | $2\sqrt{\mu_A} [\text{m}]$ |
| Characteristic of the coil pulse waveform: $\partial i_j / \partial t$ | $1[\text{A s}^{-1}]$ |

### 2.8 Formulation and solution of the inverse problem

We would like to compare inverse solutions generated with our high-resolution reciprocal solver against the widespread and well-established inverse solutions generated using MNE’s direct BEM solvers (Gramfort et al. 2013). To this end, we follow one-to-one the inverse methods implemented in MNE-Python (Gramfort et al. 2014), these methods are discussed in detail on MNE-Python’s online documentation. By doing so, we aim to eliminate any possible discrepancies due to the use of different inverse methods and ensure that different results are solely caused by the use of different modeling techniques.

We find the solution of the inverse problem as a *maximum a posteriori* solution, where given a set of prior assumptions (or distributions) on the source space, the noise, and the physical model, we find the source strength distribution of maximum likelihood —this is discussed in the Bayesian framework of Knösche and Haueisen 2022, and it is the framework used by MNE-Python (Gramfort et al. 2014; Gramfort et al. 2013). The inverse problem is then formulated using three main ingredients:

– The *M × N* leadfield matrix *L* —described in Equation 3 — which is a *modeling* term describing a linear relationship between source strengths and MEG sensor measurements;
– the source covariance matrix *R* (or source weighing matrix), which is a Bayesian *prior* term, and can be seen as an initial assumption on the properties of the source space.
– the noise covariance matrix ∑, which is an *empirical* term capturing spatiotemporal correlations of the sensor readings.

Within the leadfield matrix, we implicitly assume that the sources are constrained to the cortex, with orientation normal to the cortical surface.

In its simplest form (without source priors or noise covariance matrices), the inverse problem is formulated as a least-squares minimum norm problem (Hämäläinen and Ilmoniemi 1994), which, for an underdetermined problem (i.e., when *M < N* as is our case), can be posed as:

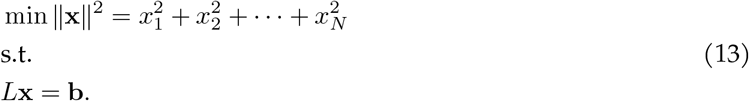

This problem has a unique solution, and the solution **x** is given explicitly by means of the pseudoinverse of *L*:

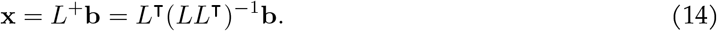

The pseudoinverse *L*^+^ = *L*^⊺^(*LL*^⊺^)^−1^ is called the *inverse operator* in this case. In general, if we want to introduce the source and noise covariance matrices, we use the following *regularized inverse operator* (Tarantola 2005; Hämäläinen and Ilmoniemi 1994; Dale and Sereno 1993):

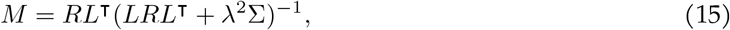

where *λ* is a regularization parameter. Note that with *R* = *I* and *λ* = 0 we recover the usual pseudoinverse. The inverse solution **x** is computed as **x** = *M* **b**. For a derivation of this regularized pseudoinverse operator, see Supplement B.

### 2.9 Depth weighting: Leadfield matrix column normalization

The sensitivity of MEG sensors to a single dipolar source depends highly on the location of the source (Hillebrand and Barnes 2002): Deeper sources produce weaker magnetic fields at the sensor positions (as predicted by the Law of Biot and Savart (Griffiths 1999)), and MEG sensors are less sensitive to the radially-oriented sources at the crests of the gyri (although these constitute a smaller fraction of the total cortical surface). Dipoles with weaker magnetic fields produce columns in the leadfield matrix *L* with a small norm; this will ultimately cause a bias towards superficial sources in the minimum norm solution.

In order to reduce this bias, we can normalize the columns of the leadfield matrix, (Fuchs et al. 1999; Lin et al. 2006). Normalizing the columns of the leadfield matrix corresponds to selecting the following source-covariance matrix:

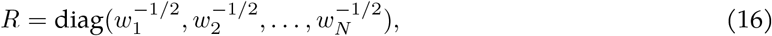

where

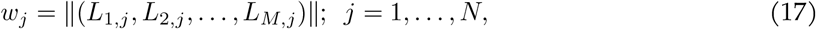

is the norm of the *j*-th column of the leadfield matrix *L*. In the MNE-Python software, we obtain this normalization by requiring the dipole orientations to be normal to the cortical surface and by setting the depth-weighing parameter *p* = 1 (Lin et al. 2006).

We also applied data whitening (Dale and Sereno 1993), which involves a transformation of the leadfield matrix given by 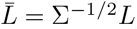,and the corresponding transformed inverse operator becomes

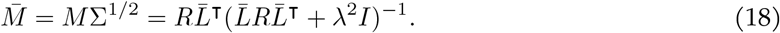

### 2.10 Dynamic Statistical Parametric Mapping (dSPM)

We can further modify our inverse solutions by employing *dynamic statistical parametric mapping (dSPM)* (Dale et al. 2000). This method utilizes a noise normalized post-hoc weighting by computing

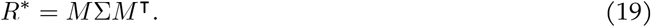

The inverse solution **x** is then normalized by

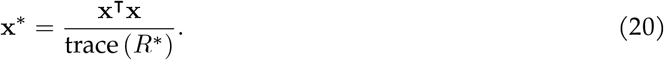

*R*^*^ is effectively the variance of the dipole strengths given a fixed dipole orientation. Therefore, if the noise covariance is computed over a sufficiently long time interval, **x**^*^ is analogous to a statistical z-score (Dale et al. 2000). That is, dSPM gives a normalized inverse solution in which the most statistically relevant dipole strengths are found among the highest percentiles. In general, the dipole strengths can vary greatly along regions of high curvature variation, i.e. along transitions from surface crests to sulci. Thus normalizing by the diagonal elements of *R*^*^ will bias the most statistically relevant solutions towards regions of least variation. Since **x** is found such that superficial sources along the surface crests are prior corrected, **x**^*^ will yield strong sources predominately in the surface sulci.

### 2.11 Evoked MEG responses to median nerve stimulation

#### Subjects, stimuli, and task

Five healthy adults participated in the recordings at the Athinoula A. Martinos Center for Biomedical Imaging, Massachusetts General Hospital (MGH), Boston, MA, between years 2004 and 2005. The study protocol was approved by the MGH Institutional Review Board (protocol #1999P010946). The subjects gave written informed consent prior to participation. The subject age range was 25–40 (mean 31 years, 2 females). Here, the subjects will be referred to by the subject codesMGH01–MGH05.

#### MRI acquisition and analysis

T1-weighted structural MRIs of the head were recorded with a magnetization-prepared rapid gradient echo (MPRAGE) sequence at 1.33 mm spatial resolution (Siemens Avanto, Trio, or Sonata; Siemens Medical Solutions, Erlangen, Germany). The MRI data were segmented using automated software and the results were used for constructing the boundary element models necessary for source modeling. For source estimation with BEM-FMM, the MRI data were segmented using the `headreco` pipeline of SimNIBS (Nielsen et al. 2018). For source estimation using MNE the MRI data were segmented and 3-layer BEMs (scalp, outer skull, inner skull) built using the Freesurfer software (Reuter et al. 2012).

#### MEG recordings and sensor-space analysis

Whole-head 306-channel MEG (VectorView; MEGIN Oy, Finland) data were recorded in a magnetically shielded room (Cohen et al. 2002). The instrument employs sensor triplets (one magnetometer and two planar gradiometers) at 102 measurement locations. Simultaneously, 70-channel EEG (not analyzed here) as well as horizontal and vertical electrooculogram (EOG) were recorded. All signals were bandpass-filtered to 0.01–250 Hz and sampled at 1 kHz.

At the start of the MEG session, the locations of 4 head position indicator (HPI) coils attached to the scalp and several additional scalp surface points were recorded with respect to fiduciary landmarks (nasion and two preauricular points) using a 3-D digitizer (Fastrak Polhemus, VT, USA). During MRI–MEG coordinate system alignment, the fiduciary points were then identified from the structural MRIs. Using the HPI and scalp surface locations, this initial approximation was refined using an iterative closest-point search algorithm.

During the MEG recordings, the subjects received 0.2 ms square wave pulses at a suprathreshold intensity at the right wrist over the median nerve (Telefactor S88, Grass Instrument Company, Quincy, MA, USA). The stimuli were presented at a pseudorandom interstimulus interval of 3–9 seconds. The task was to respond to each stimulus by lifting the left hand index finger as quickly as possible.

Evoked responses were averaged with respect to the median nerve stimuli in a time window from 200 ms prestimulus to 1150 ms poststimulus with a band-pass of 0.3–1000 kHz. Epochs containing EOG signals exceeding 150 mV peak-to-peak amplitude were automatically discarded from the averages. After rejecting trials containing artifacts, the evoked responses contained 79 *±*19 (mean *±*standard deviation) trials. The noise covariance matrix was estimated from a time window of -200 to -20 ms relative to the median nerve stimuli. Figure 2A shows an example of the evoked gradiometer signals and sensor topographies for subject MGH01.

**Figure 2.**
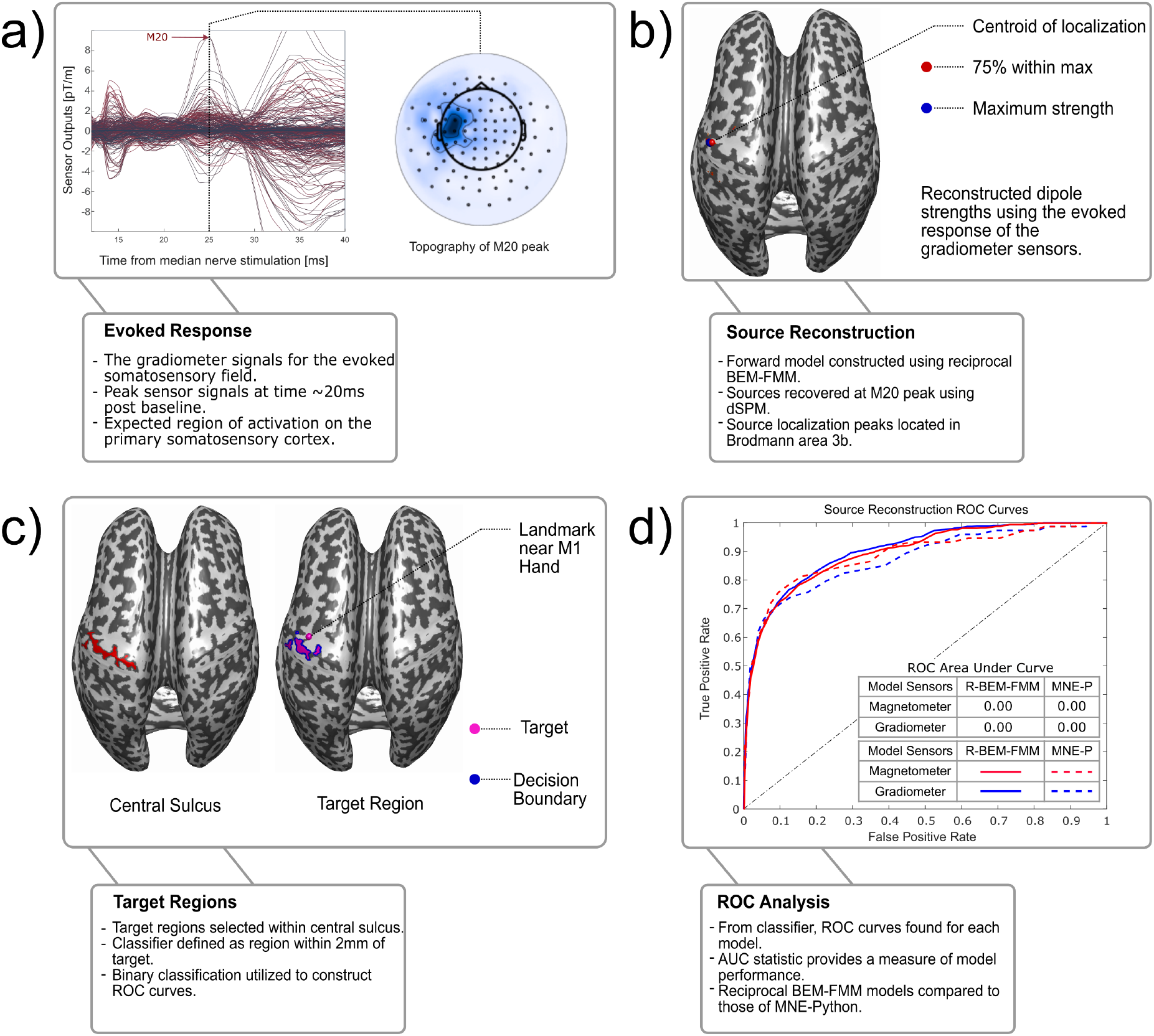
The workflow of the source localization ROC analysis, using subject MGH01. a) The gradiometer signals for the evoked somatosensory field, along with their corresponding topographies. The M20 peak occurs at a latency of approximately 20ms post stimulation. b) The Reciprocal BEM-FMM source localization dSPM solution for M20 peak plotted on the inflated white matter. The centroid of 75% strength threshold is plotted as a red sphere, and the maximum strength is plotted as a blue sphere. c) Plot of the tagged central sulcus (left,red), the approximate M1 hand (right, magenta sphere), and the tagged true-positive region for binary classification (right, magenta) and *decision boundary* (right,blue), i.e. the boundary of the true-positive region. d) The ROC curves for the reciprocal BEM-FMM (solid line) and MNE-Python (dashed line) source localization models. We compare models generated from gradiometer (blue) and magnetometer (red) data. AUC values for these models can be found in Table 3.

For the MNE-Python source localizations, the source space was generated with the `setup_source_space` procedure with `ico5` source spacing, which produces 20477 *±* 5 source dipoles evenly spread across the cortex. In our BEM-FMM source localization, the source space consists of the triangle centers of the white matter mesh of each subject. The tissue layers, together with their conductivity values, average number of triangles, and relevant bibliographic references for both our BEM-FMM models and the MNE-Python models are included in Table 2, see also (SimNIBS Conductivities 2025).

**Table 2:** Conductivities and number of triangles in each tissue layer for the reciprocal BEM-FMM and MNE-Python models.

| Tissue | Conductivity [$\text{S m}^{-1}$] | avg. tri. | Reference |
| --- | --- | --- | --- |
| Skin | 0.25 | 231 000 | (Truong et al. 2012) |
| Skull | 0.01 | 237 000 | (Wagner et al. 2004) |
| CSF | 1.654 | 126 000 | (Wagner et al. 2004) |
| GM | 0.275 | 347 000 | (Wagner et al. 2004; Thielscher and Wichmann 2009) |
| WM | 0.126 | 384 000 | (Wagner et al. 2004; Thielscher and Wichmann 2009) |
| Ventricles | 1.654 | 14 800 | (Wagner et al. 2004)<sup>†</sup> |
| Eyes | 0.5 | 5070 | (Gabriel et al. 1996; Opitz et al. 2015) |

| Tissue name | Conductivity [$\text{S m}^{-1}$] | Reference |
| --- | --- | --- |
| Skin | 0.25 | (Truong et al. 2012) |
| Outer Skull | 0.01 | (Wagner et al. 2004) |
| Inner Skull | 0.33 | (Vorwerk et al. 2019) |

(b) Tissue layers in the MNE-Python inverse models and their conductivities. Each layer contains 20480 triangular faces.

**Table 3:** The AUC values of the source localization results for every subject model for both reciprocal BEM-FMM and MNE-Python. AUC values closer to 1 indicate a better model performance.

(a) AUC values for source localization using only the magnetometer recordings at the M20 peak.
| Magnetometer Sensors | MGH01 | MGH02 | MGH03 | MGH04 | MGH05 | Mean |
| --- | --- | --- | --- | --- | --- | --- |
| R-BEM-FMM | 0.8972 | 0.8113 | 0.8414 | 0.9365 | 0.9265 | 0.8826 |
| MNE-Python | 0.8863 | 0.7692 | 0.7802 | 0.8026 | 0.8326 | 0.8142 |

(b) AUC values for source localization using only the gradiometer recordings at the M20 peak.
| Gradiometer Sensors | MGH01 | MGH02 | MGH03 | MGH04 | MGH05 | Mean |
| --- | --- | --- | --- | --- | --- | --- |
| R-BEM-FMM | 0.9033 | 0.7614 | 0.8250 | 0.9556 | 0.9374 | 0.8765 |
| MNE-Python | 0.8723 | 0.7906 | 0.7973 | 0.7694 | 0.8581 | 0.8175 |

### 2.12 Visualization of the reconstructed dipole strength densities

We visualized the normalized reconstructed source strength density exceeding a specified threshold *T* relative to the maximum strength. Specifically, if *s*_max_ represents the largest source strength magnitude, then we plot all sources with strength *s* ≥ *T · s*_max_. Additionally, we plot the centroid (e.g. geometric mean) of the region with sources exceeding a given threshold. Figure 2B shows an example of the centroid at a 75% threshold (as well as the source of maximum strength) of the reciprocal BEM-FMM dSPM solution for subject MGH01.

To compare the results of MNE-Python’s and BEM-FMM source reconstructions, we plotted the reconstructed dipole source density on the same inflated white matter mesh (Fischl et al. 1999) obtained from the subject `headreco` segmentations. Whenever coregistration was needed, we obtained optimal transformations using ICP. The source strength densities obtained from MNE-Python are sparse relative to the mesh, and hence were smoothed iteratively to replicate the native visualization of MNE-Python. The smoothing is performed with a *k*-nearest neighbors algorithm at each triangle center, and averaging the neighbor values.

### 2.13 The Statistical ROC metric

The *receiver operating characteristic (ROC)* curve is a tool to evaluate the performance of binary classification models (Knösche and Haueisen 2022; Hastie et al. 2009). We treat source localization as a binary classification problem by specifying a target region for the “true” dipole sources. To avoid adding biases, we selected a sufficiently large subregion of the central sulcus closest to the M1-hand region, which we tagged manually. This is done computationally by 1) tagging the mesh triangles of the white matter that lie in central sulcus; and 2) choosing a subregion of the tagged central sulcus, consisting of 40% of the total central sulcus area closest to the M1-hand region.

The tagging of the central sulcus points is done by manually selecting points along the sulcus and then using a *k*-nearest neighbors search to locate the nearest mesh faces. Among this set of faces, we tag only those with a mean curvature value less than 1 (mean curvature *H <* 1 indicates the presence of a “valley” on the surface (do Carmo 2016)). From a selected point in the M1-hand region, we chose the smallest radius *R* such that the area of the triangles with centers at distance *< R* from the M1-hand point is at least 40% of the total area of the central sulcus tagged. These flagged points near the M1-hand are our target region of true sources for binary classification. The central sulcus and target region tagged for the subject MGH01 are seen in Figure 2C.

We classify sources as *true positives* (TP), *false negatives* (FN), *false positives* (FP), and *true negatives* (TN), based on the following procedure: begin with a given threshold percentage *T* ; if a source dipole on the target region has a normalized source strength *s > T*, then the source dipole is considered a TP; similarly, if *s < T*, then it is a FN. Likewise, if a source dipole outside the target region has a normalized source strength *s > T*, then it is a FP; if *s < T*, then it is a TN.

With this binary classification, the ROC curve is generated by plotting the rate of false positives, *FP/*(*FP* + *TN*), versus the rate of true positives, *TP/*(*TP* + *FN*), as they vary across all threshold percentages *T* from 0 to 100%. The *area under the ROC curve (AUC)* serves as a numerical measure of the efficacy of the model: if *AUC* = 1, then the model is a perfect classifier; whereas if *AUC* ≤ 0.5, then the model behaves no better than a random binary classifier (Hastie et al. 2009). Example ROC curves for subject MGH01 are found in Figure 2D.

### 2.14 Model evaluation with simulated ground truth data

#### Direct simulation of evoked somatosensory fields from equivalent current dipoles

For each of the 5 subjects, we manually selected a location for a tangentially-oriented dipole on the posterior wall of the central sulcus (Brodmann area 3, see Figure 3).

**Figure 3.**
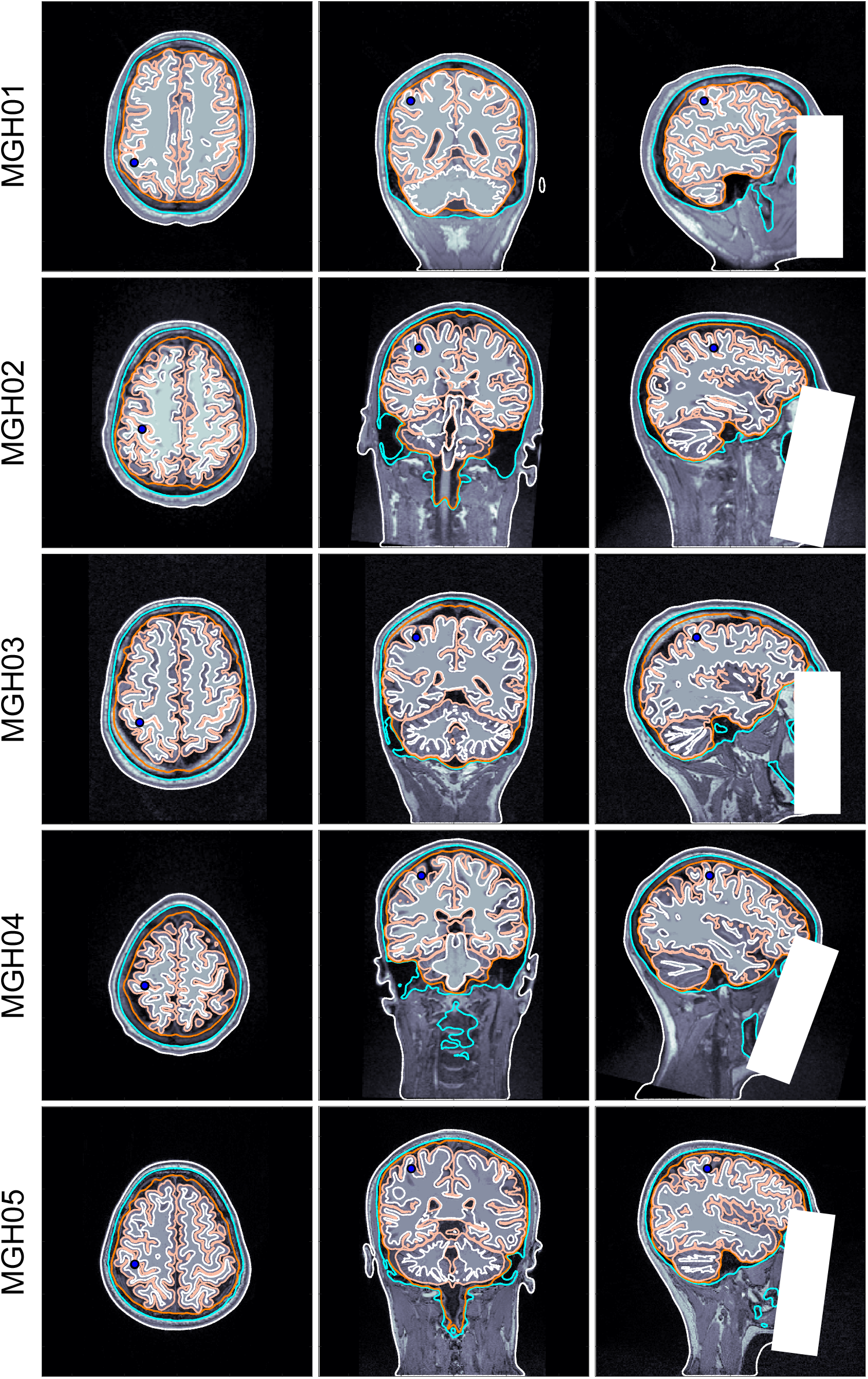
Placement of selected dipoles (blue circles) on Brodmann area 3b for each of the 5 subjects. Each tissue layer used in the model has the following color assignation: Skin —white (outer); Skull —cyan; CSF —orange; GM —apricot; WM —white (inner).

The **B**-field corresponding to each selected dipole was generated using BEM-FMM with *b*-refinement (Makarov et al. 2021; Wartman et al. 2025), and sampled at 64 observation points for each magnetometer sensor, and 128 observation points for each gradiometer sensor. The sensor readings were then computed from the values of the flux **B** at the observation points by approximating the surface integral

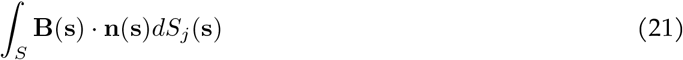

on the surface *S*_*j*_ enclosed by sensor *j* (Knösche and Haueisen 2022). In the case of magnetometers, this integral is approximated directly, see Figure 4. In the case of gradiometer pairs, the normal vectors of each coil are taken in opposite directions and the integral is divided by the distance separating each coil. The simulated magnetometer and gradiometer signals were used for source reconstruction to evaluate the performance of our inverse method in the absence of noise.

**Figure 4.**
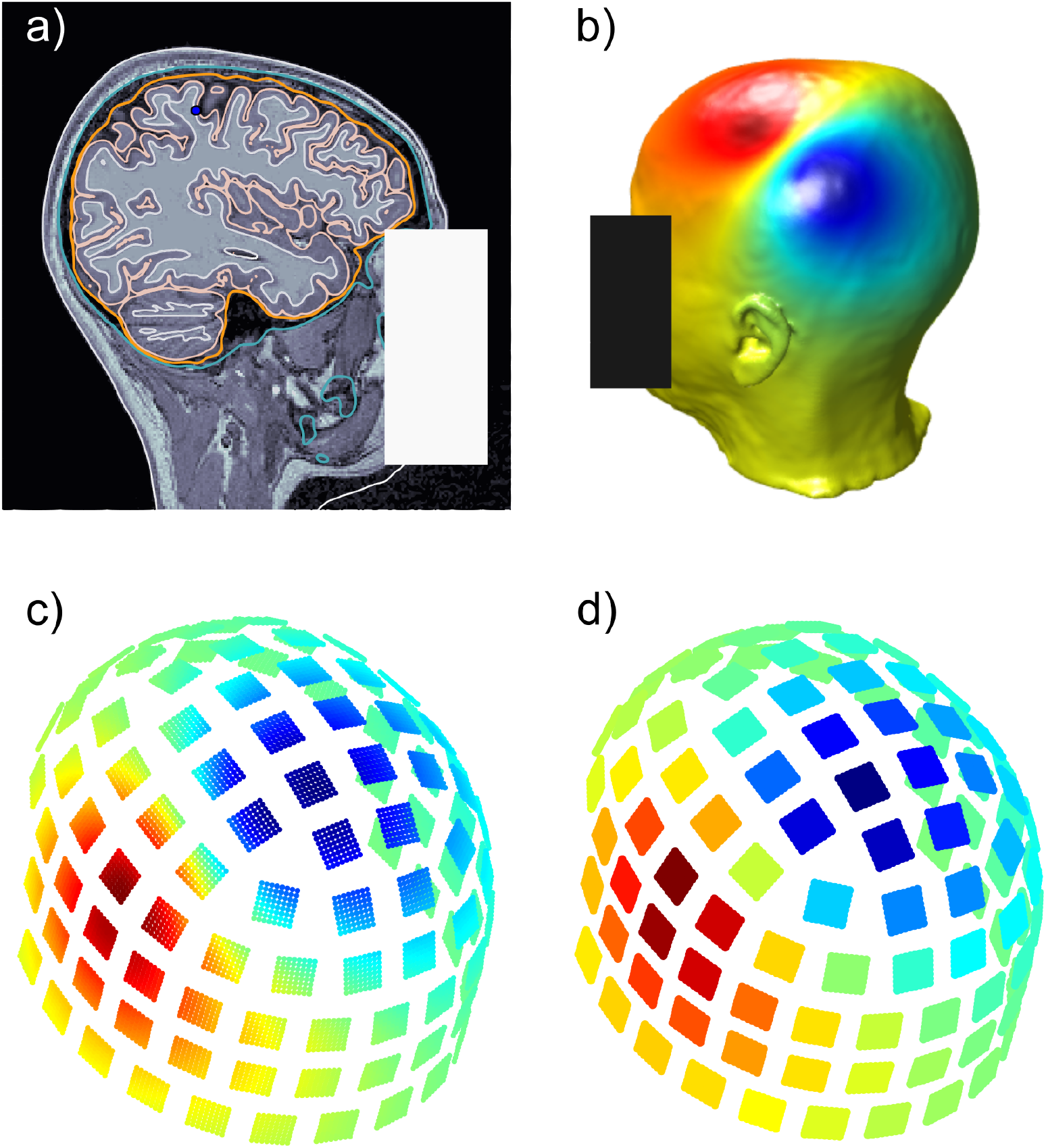
Workflow for the generation of simulated magnetometer data: a) dipole placement in the primary somatosensory cortex for subject MGH05; b) surface potential on the skin calculated BEM-FMM with *b*-refinement; c) **B**-field calculated at 64 observation points for each magnetometer; d) simulated sensor outputs by numerical computation of the integral of the normal component of **B** at the observation points.

#### Source localization of simulated noisy somatosensory evoked fields at various SNR levels

We tested the stability of our inverse method by incorporating different levels of noise to our simulated MEG signals corresponding to the selected dipoles. The noise was modeled by randomly sampling 2000 dipoles across the entire cortical surface: each dipole was centered on the midsurface between the grey and white matter surfaces and oriented normally to the GM surface. For each of the randomly selected dipoles, we computed the forward BEM-FMM solution with *b*-AMR and created a direct leadfield matrix *L* of size 306 *×* 2000 by letting each column be the MEG sensor signals corresponding to each dipole. We generated 1000 samples of noisy signals at each target signal-to-noise ratio level with the following procedure:

1. For each *ω* = 1, …, 1000 we sampled a random 2000 *×* 1 vector of dipole strengths **X**_*ω*_ ∼ 𝒢 (0, *σ*) following a gaussian distribution of zero mean and standard deviation *σ* = 1.
2. By the quasi-static Maxwell assumption, the noise ***ϵ***_*ω*_ is computed from the direct leadfield matrix as

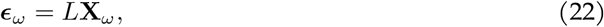

and it is then divided by its norm, so that ***ϵ***_*ω*_ is a unit vector.
3. The total signal was defined in terms of a parameter *λ* between 0 and 1 as

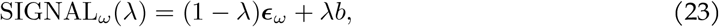

where *b* is the normalized (unit vector) MEG signals of the selected dipole in the posterior wall of the central sulcus (described above). The signal is subsequently re-scaled to a realistic magnitude (Murakami and Okada 2015).
4. The SNR is calculated according to the formula:

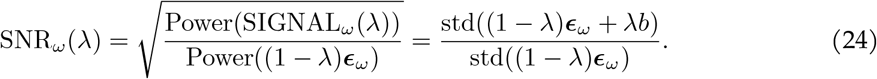

A value *λ*^*^ between 0 and 1 was calculated numerically, so that

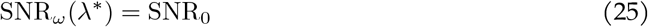

for each target signal-to-noise ration SNR_0_.

The SNR values we tested were 81, 27, 3, 2, and 1.5. For each sample at every SNR level, we performed source localization and recorded the distance from the source dipole to 1) the reconstructed source of maximum strength (peak) and 2) to the centroid of the 75% activation threshold.

Both for the noiseless source localization, and the source localization at different SNR levels, we employed the minimum norm estimation described in Section 2.8, without the use of dSPM. The reason for the omission of the dSPM step is that this method requires the estimation of a noise covariance matrix, which for simulated data may result in the introduction of biases.

#### Error maps for source localization at various noise levels

Using the forward BEM-FMM solution for each of the 2000 dipoles placed uniformly at random over the cortex —which we described above— we generated error maps indicating the average localization error at several *absolute noise levels* at each of the dipole locations. For each dipole, of index *j*, we created 500 samples of noise signals by multiplying the 306 *×* 2000 lead-field matrix *L* by a random vector of strengths **X**_*ω*_, where we impose the condition that **X**_*ω*_(*j*) = 0, so that the noise does not contribute to the source signal. The total signal was computed for a *fixed* absolute noise percentage 0 ≤ *λ*_0_ ≤ 1 as

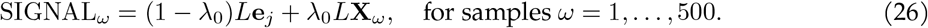

Here, **e**_*j*_ is the vector with all entries equal to 0 except at the position *j* where it has the value 1. For each of the sample signals, we performed source reconstruction and measured the distance from the centroid of the estimated sources at the 75% thereshold to the source dipole, as well as the distance from the peak of the estimated region of activation to the source. The reason for taking a fixed absolute noise level rather than a fixed SNR is that having a fixed SNR will force deep sources to have a much larger magnitude than the superficial ones. So using a fixed SNR limits the comparability of the quality of fits among different cortical regions and different sensor data. By keeping a fixed noise level, we avoid these issues.

## 3 Results

### Experimental data: evoked MEG responses to median nerve stimulation

Figures 5 and 6 show the source localization peaks, centroids and 50% threshold activation for the evoked somatosensory fields for each subject calculated using dSPM. Each figure shows the solutions computed using reciprocal BEM-FMM and MNE-Python, with solutions in Figure 5 computed from gradiometer data, and solutions in Figure 6 computed from magnetometer data. We computed ROC curves from each of these models, which can be found in Supplement C. Table 3 shows the AUC values corresponding to each subject model using both reciprocal BEM-FMM and MNE-Python.

**Figure 5.**
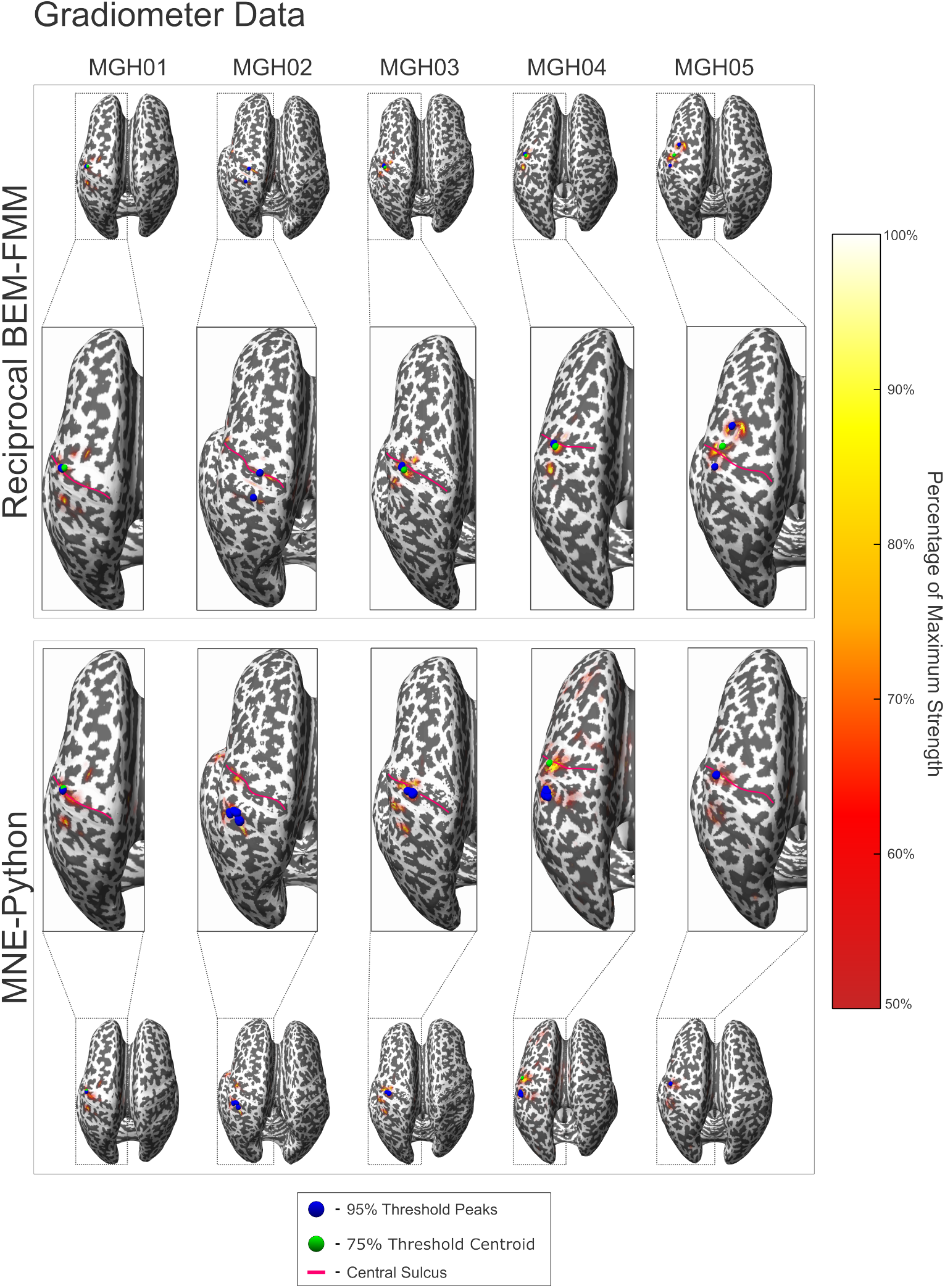
Reciprocal BEM-FMM and MNE-Python source localization results for all subjects using gradiometer data and dSPM. The blue spheres indicate sources with strengths within 95% of the maximum (peaks). The green sphere indicates the centroid of all localized sources with strengths within 75% of the maximum. The red curves denote the central sulcus of each subject. The colorbar denotes sources with strengths within 50-100% of the maximum.

**Figure 6.**
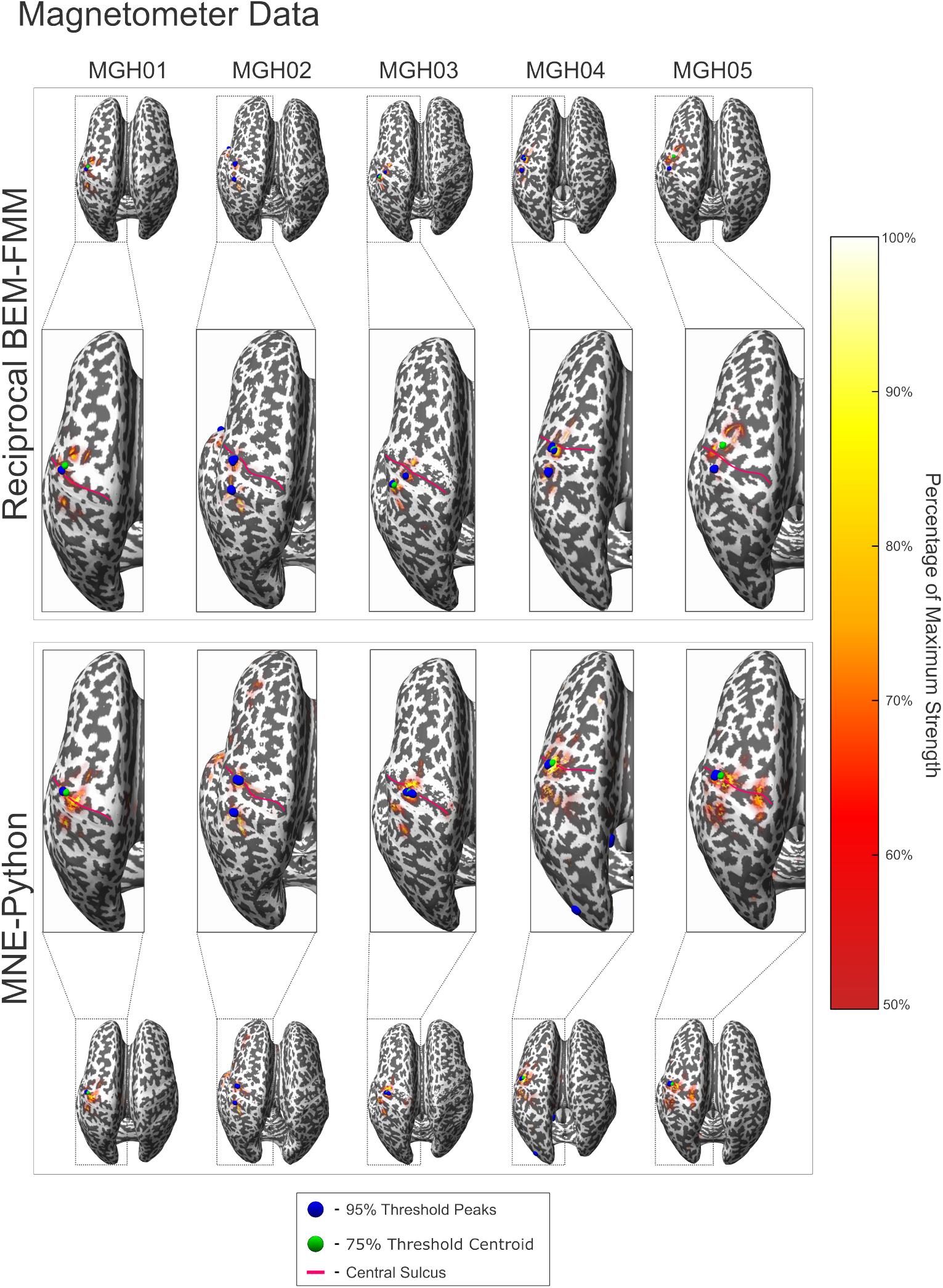
Reciprocal BEM-FMM and MNE-Python source localization results for all subjects using magnetometer data and dSPM. The blue spheres indicate sources with strengths within 95% of the maximum (peaks). The green sphere indicates the centroid of all localized sources with strengths within 75% of the maximum. The red curves denote the central sulcus of each subject. The colorbar denotes sources with strengths within 50-100% of the maximum.

### Synthetic data: source reconstruction for simulated somatosensory fields at various noise levels

Figures 8 and 7 show the source localization peaks, centroids, and 50% threshold activation for the noiseless evoked somatosensory fields of each subject, where noiseless synthetic MEG signals were generated from the indicated source dipole. Tables 4a and 4b show the aggregated distance error statistics corresponding to the centroid-to-source and peak-to-source distances, respectively, from the ground truth source across varying levels of noise, using synthetic magnetometer and gradiometer data. Values in Table 4 are averaged across each subject; complete statistical data tables for individual subjects are shown in Supplement C.

**Table 4:** Aggregated distance error statistics for source estimation of simulated data (at various noise levels) for a dipole placed in the primary somatosensory cortex: The distance from centroids of the estimated region of activation, and from the peak of activation, to the true source location are computed and averaged for each subject.

| SNR | Magnetometer Data |  |  | Gradiometer Data |  |  |
| --- | --- | --- | --- | --- | --- | --- |
|  | Mean | Median | Std | Mean | Median | Std |
| Noiseless | 6.67 | - | - | 6.09 | - | - |
| 81 | 6.49 | 6.52 | 0.05 | 6.71 | 6.67 | 0.07 |
| 27 | 6.50 | 6.46 | 0.11 | 6.74 | 6.67 | 0.13 |
| 9 | 6.51 | 6.42 | 0.18 | 6.77 | 6.71 | 0.22 |
| 3 | 6.76 | 6.51 | 0.75 | 6.71 | 6.79 | 0.58 |
| 2 | 7.17 | 6.78 | 1.19 | 6.80 | 6.83 | 0.85 |
| 1.5 | 7.90 | 6.20 | 3.40 | 7.08 | 5.94 | 5.58 |

| SNR | Magnetometer Data |  |  | Gradiometer Data |  |  |
| --- | --- | --- | --- | --- | --- | --- |
|  | Mean | Median | Std | Mean | Median | Std |
| Noiseless | 10.08 | - | - | 9.92 | - | - |
| 81 | 10.38 | 10.38 | 0.00 | 9.54 | 9.75 | 0.22 |
| 27 | 10.38 | 10.38 | 0.03 | 9.53 | 9.31 | 0.24 |
| 9 | 10.44 | 10.38 | 0.30 | 9.58 | 9.45 | 0.37 |
| 3 | 10.72 | 10.38 | 1.63 | 9.64 | 9.54 | 0.47 |
| 2 | 11.32 | 10.44 | 2.55 | 9.68 | 9.48 | 0.80 |
| 1.5 | 12.31 | 10.56 | 6.14 | 11.63 | 9.64 | 10.17 |

**Table 5:** Statistical results for centroid-to-source distance error with noiseless and noisy simulated data for every MGH subject. Centroids are taken from the set of sources with strengths within 75-100% of the maximum strength.

| MGH01 |  |  |  |  |  |  |
| --- | --- | --- | --- | --- | --- | --- |
| SNR | Magnetometer Data |  |  | Gradiometer Data |  |  |
|  | Mean | Median | Std | Mean | Median | Std |
| Noiseless | 4.28 | - | - | 4.34 | - | - |
| 81 | 2.93 | 3.07 | 0.15 | 6.03 | 5.97 | 0.07 |
| 27 | 2.88 | 2.77 | 0.14 | 6.03 | 5.97 | 0.07 |
| 9 | 2.91 | 2.77 | 0.18 | 6.03 | 5.97 | 0.15 |
| 3 | 3.73 | 3.07 | 1.17 | 5.50 | 5.97 | 1.10 |
| 2 | 4.32 | 3.71 | 1.40 | 5.45 | 5.97 | 1.17 |
| 1.5 | 5.62 | 4.91 | 6.04 | 8.33 | 6.00 | 10.38 |

| MGH02 |  |  |  |  |  |  |
| --- | --- | --- | --- | --- | --- | --- |
| SNR | Magnetometer Data |  |  | Gradiometer Data |  |  |
|  | Mean | Median | Std | Mean | Median | Std |
| Noiseless | 12.50 | - | - | 11.01 | - | - |
| 81 | 13.12 | 13.12 | 0.03 | 11.27 | 11.16 | 0.22 |
| 27 | 13.06 | 13.12 | 0.10 | 11.34 | 11.16 | 0.31 |
| 9 | 12.93 | 12.94 | 0.28 | 11.32 | 11.16 | 0.42 |
| 3 | 12.75 | 12.86 | 0.61 | 11.25 | 11.16 | 0.73 |
| 2 | 12.94 | 12.88 | 1.21 | 11.30 | 11.20 | 1.11 |
| 1.5 | 13.47 | 12.94 | 2.61 | 11.85 | 11.63 | 2.99 |

| MGH03 |  |  |  |  |  |  |
| --- | --- | --- | --- | --- | --- | --- |
| SNR | Magnetometer Data |  |  | Gradiometer Data |  |  |
|  | Mean | Median | Std | Mean | Median | Std |
| Noiseless | 6.09 | - | - | 5.89 | - | - |
| 81 | 5.69 | 5.68 | 0.04 | 5.32 | 5.32 | 0.00 |
| 27 | 5.74 | 5.68 | 0.15 | 5.32 | 5.32 | 0.01 |
| 9 | 5.82 | 5.68 | 0.21 | 5.29 | 5.32 | 0.15 |
| 3 | 5.87 | 5.76 | 0.24 | 5.28 | 5.32 | 0.27 |
| 2 | 5.90 | 5.98 | 0.41 | 5.41 | 5.32 | 0.44 |
| 1.5 | 6.17 | 5.98 | 4.10 | 6.85 | 5.68 | 6.26 |

| MGH04 |  |  |  |  |  |  |
| --- | --- | --- | --- | --- | --- | --- |
| SNR | Magnetometer Data |  |  | Gradiometer Data |  |  |
|  | Mean | Median | Std | Mean | Median | Std |
| Noiseless | 5.00 | - | - | 3.97 | - | - |
| 81 | 4.89 | 4.88 | 0.04 | 4.79 | 4.75 | 0.07 |
| 27 | 4.97 | 4.88 | 0.14 | 4.82 | 4.75 | 0.09 |
| 9 | 5.03 | 4.88 | 0.16 | 4.85 | 4.93 | 0.10 |
| 3 | 5.38 | 4.93 | 1.35 | 4.95 | 4.93 | 0.18 |
| 2 | 6.37 | 5.19 | 2.33 | 5.02 | 4.93 | 0.62 |
| 1.5 | 7.49 | 5.75 | 2.84 | 6.92 | 4.98 | 6.83 |

| MGH05 |  |  |  |  |  |  |
| --- | --- | --- | --- | --- | --- | --- |
| SNR | Magnetometer Data |  |  | Gradiometer Data |  |  |
|  | Mean | Median | Std | Mean | Median | Std |
| Noiseless | 5.50 | - | - | 5.25 | - | - |
| 81 | 5.83 | 5.83 | 0.00 | 6.17 | 6.17 | 0.00 |
| 27 | 5.83 | 5.83 | 0.00 | 6.22 | 6.17 | 0.16 |
| 9 | 5.85 | 5.83 | 0.10 | 6.36 | 6.17 | 0.30 |
| 3 | 6.07 | 5.94 | 0.38 | 6.57 | 6.55 | 0.64 |
| 2 | 6.31 | 6.15 | 0.62 | 6.80 | 6.72 | 0.93 |
| 1.5 | 6.74 | 1.44 | 1.44 | 1.44 | 1.44 | 1.44 |

**Table 6:** Statistical results for peak-to-source distance error with noiseless and noisy simulated data for every MGH subject.

| MGH01 |  |  |  |  |  |  |
| --- | --- | --- | --- | --- | --- | --- |
| SNR | Magnetometer Data |  |  | Gradiometer Data |  |  |
|  | Mean | Median | Std | Mean | Median | Std |
| Noiseless | 7.43 | - | - | 9.54 | - | - |
| 81 | 7.43 | 7.43 | 0.00 | 9.54 | 9.54 | 0.00 |
| 27 | 7.46 | 7.43 | 0.13 | 9.54 | 9.54 | 0.00 |
| 9 | 7.58 | 7.43 | 0.26 | 9.51 | 9.54 | 0.08 |
| 3 | 7.71 | 7.43 | 0.30 | 9.41 | 9.54 | 0.19 |
| 2 | 7.79 | 8.00 | 0.32 | 9.26 | 9.25 | 0.34 |
| 1.5 | 8.67 | 8.00 | 7.02 | 12.60 | 9.25 | 15.87 |

| MGH02 |  |  |  |  |  |  |
| --- | --- | --- | --- | --- | --- | --- |
| SNR | Magnetometer Data |  |  | Gradiometer Data |  |  |
|  | Mean | Median | Std | Mean | Median | Std |
| Noiseless | 16.45 | - | - | 13.27 | - | - |
| 81 | 16.45 | 16.45 | 0.00 | 13.27 | 13.27 | 0.00 |
| 27 | 16.45 | 16.45 | 0.00 | 13.28 | 13.27 | 0.12 |
| 9 | 16.62 | 16.45 | 1.12 | 13.53 | 13.27 | 0.45 |
| 3 | 18.01 | 16.45 | 3.01 | 13.65 | 13.27 | 0.60 |
| 2 | 18.51 | 16.45 | 3.42 | 13.79 | 13.27 | 1.74 |
| 1.5 | 18.99 | 16.58 | 5.07 | 15.26 | 14.06 | 7.62 |

| MGH03 |  |  |  |  |  |  |
| --- | --- | --- | --- | --- | --- | --- |
| SNR | Magnetometer Data |  |  | Gradiometer Data |  |  |
|  | Mean | Median | Std | Mean | Median | Std |
| Noiseless | 11.12 | - | - | 8.43 | - | - |
| 81 | 11.12 | 11.12 | 0.00 | 8.43 | 8.43 | 0.00 |
| 27 | 11.12 | 11.12 | 0.00 | 8.43 | 8.43 | 0.00 |
| 9 | 11.12 | 11.12 | 0.01 | 8.43 | 8.43 | 0.00 |
| 3 | 10.75 | 11.12 | 0.76 | 8.44 | 8.43 | 0.07 |
| 2 | 10.37 | 10.85 | 1.06 | 8.49 | 8.43 | 0.20 |
| 1.5 | 10.68 | 10.41 | 7.78 | 10.01 | 8.43 | 10.48 |

| MGH04 |  |  |  |  |  |  |
| --- | --- | --- | --- | --- | --- | --- |
| SNR | Magnetometer Data |  |  | Gradiometer Data |  |  |
|  | Mean | Median | Std | Mean | Median | Std |
| Noiseless | 6.69 | - | - | 7.59 | - | - |
| 81 | 6.09 | 6.09 | 0.00 | 6.55 | 7.59 | 1.09 |
| 27 | 6.09 | 6.09 | 0.00 | 6.48 | 5.42 | 1.09 |
| 9 | 6.09 | 6.09 | 0.00 | 6.43 | 6.09 | 1.05 |
| 3 | 6.75 | 6.09 | 3.33 | 6.47 | 6.53 | 1.03 |
| 2 | 9.73 | 6.09 | 7.12 | 6.52 | 6.53 | 1.24 |
| 1.5 | 13.02 | 7.59 | 8.77 | 9.94 | 6.53 | 16.39 |

| MGH05 |  |  |  |  |  |  |
| --- | --- | --- | --- | --- | --- | --- |
| SNR | Magnetometer Data |  |  | Gradiometer Data |  |  |
|  | Mean | Median | Std | Mean | Median | Std |
| Noiseless | 8.70 | - | - | 10.76 | - | - |
| 81 | 10.79 | 10.79 | 0.00 | 9.92 | 9.92 | 0.00 |
| 27 | 10.79 | 10.79 | 0.00 | 9.92 | 9.92 | 0.00 |
| 9 | 10.79 | 10.79 | 0.10 | 10.00 | 9.92 | 0.26 |
| 3 | 10.40 | 10.79 | 0.73 | 10.26 | 9.92 | 0.45 |
| 2 | 10.18 | 10.79 | 0.85 | 10.33 | 9.92 | 0.47 |
| 1.5 | 10.21 | 10.20 | 2.06 | 10.32 | 9.92 | 0.51 |

**Figure 7.**
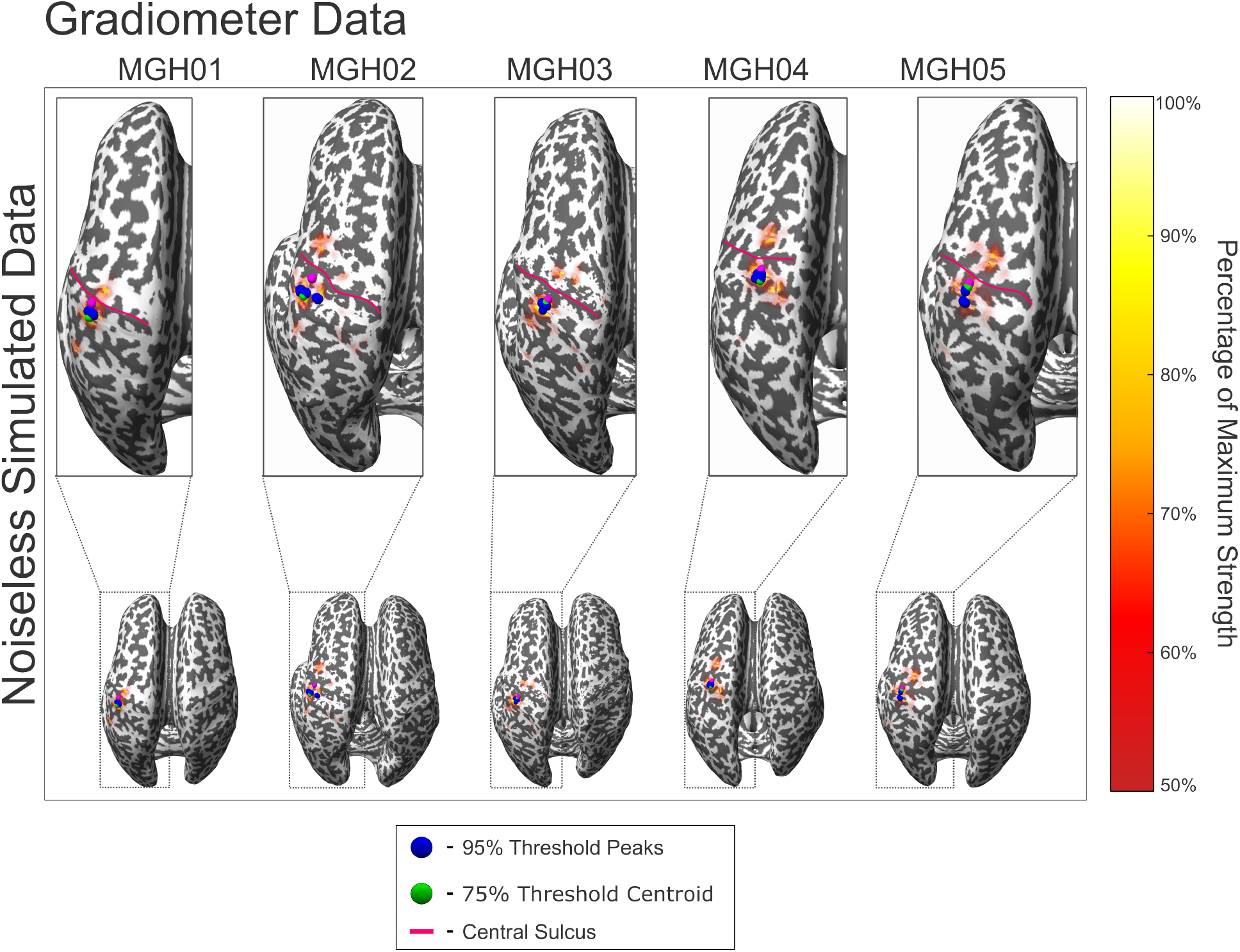
Reciprocal BEM-FMM source localization results for all subjects using synthetic gradiometer signals without noise. The magenta sphere indicates the source dipole used to generate the synthetic data. The blue spheres indicate sources with strengths within 95% of the maximum (peaks). The green sphere indicates the centroid of all localized sources with strengths within 75% of the maximum. The red curves denote the central sulcus of each subject. The colorbar denotes sources with strengths within 50-100% of the maximum.

**Figure 8.**
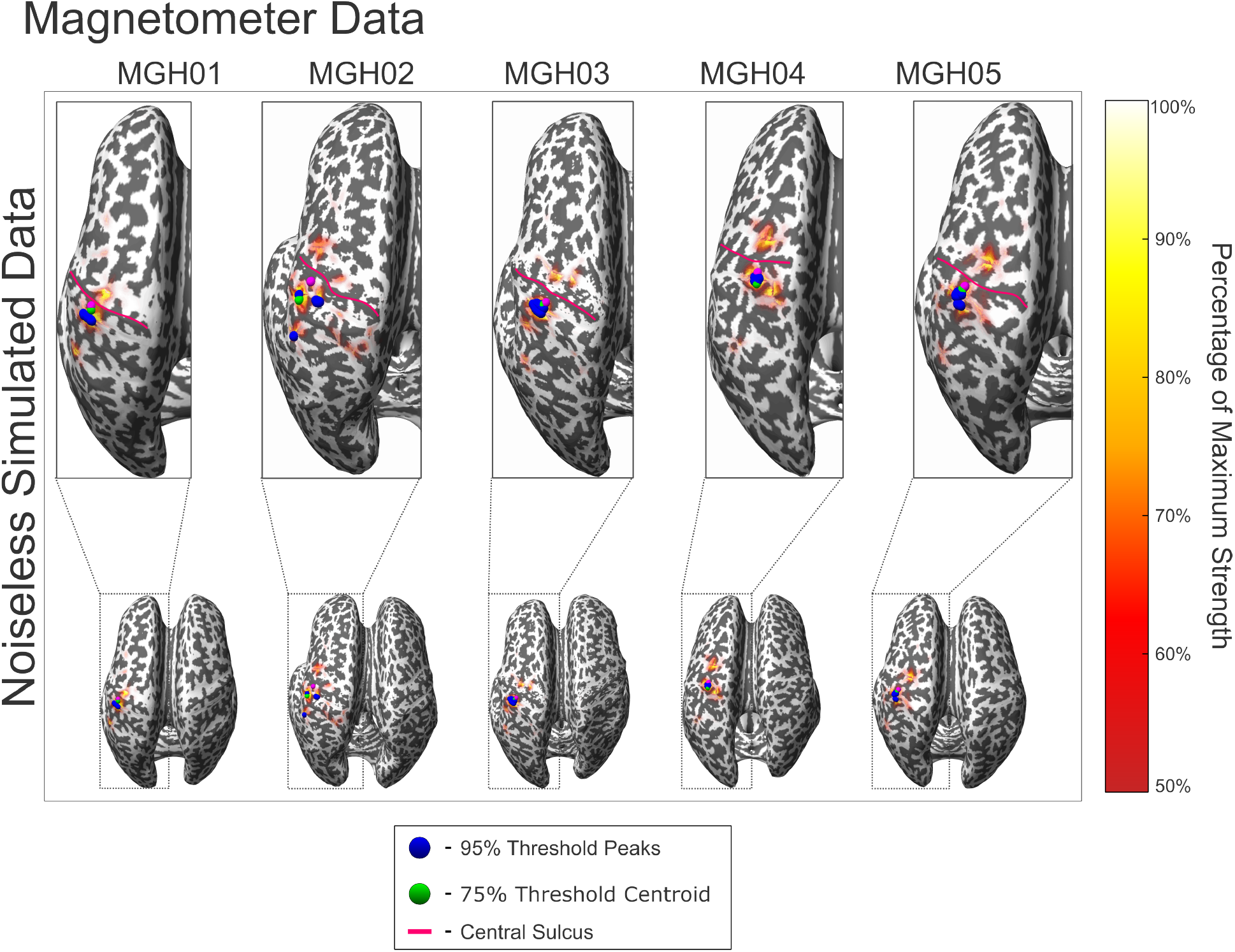
Reciprocal BEM-FMM source localization results for all subjects using synthetic noiseless magnetometer signals. The magenta sphere indicates the source dipole used to generate the synthetic data. The blue spheres indicate sources with strengths within 95% of the maximum (peaks). The green sphere indicates the centroid of all localized sources with strengths within 75% of the maximum. The red curves denote the central sulcus of each subject. The colorbar denotes sources with strengths within 50-100% of the maximum.

Figure 9 shows the source localization error maps for subject MGH01.

**Figure 9.**
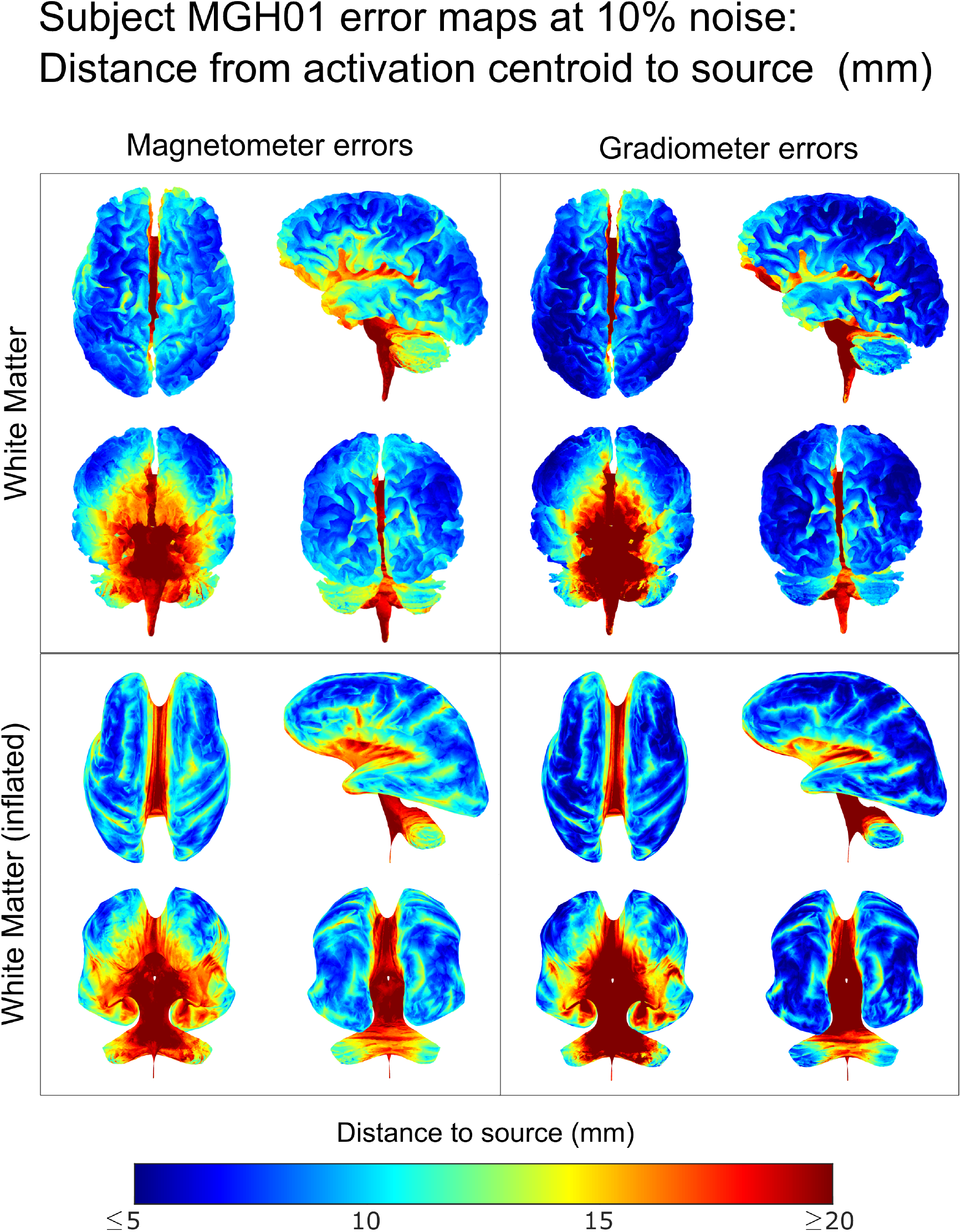
Error maps at 10% noise level (mm) for subject MGH01 on the white matter and inflated white matter surfaces. At each surface point, the average distance from the source to the centroid of estimated source strengths is interpolated from the exact computation of 2000 dipole locations chosen uniformly at random with 500 samples of noise at 10% level for each dipole location. In each box, we display different views of each tissue: top left — dorsal transverse view; top right — left lateral sagittal view; bottom left — anterior coronal view; bottom right — posterior coronal view.

**Figure 10.**
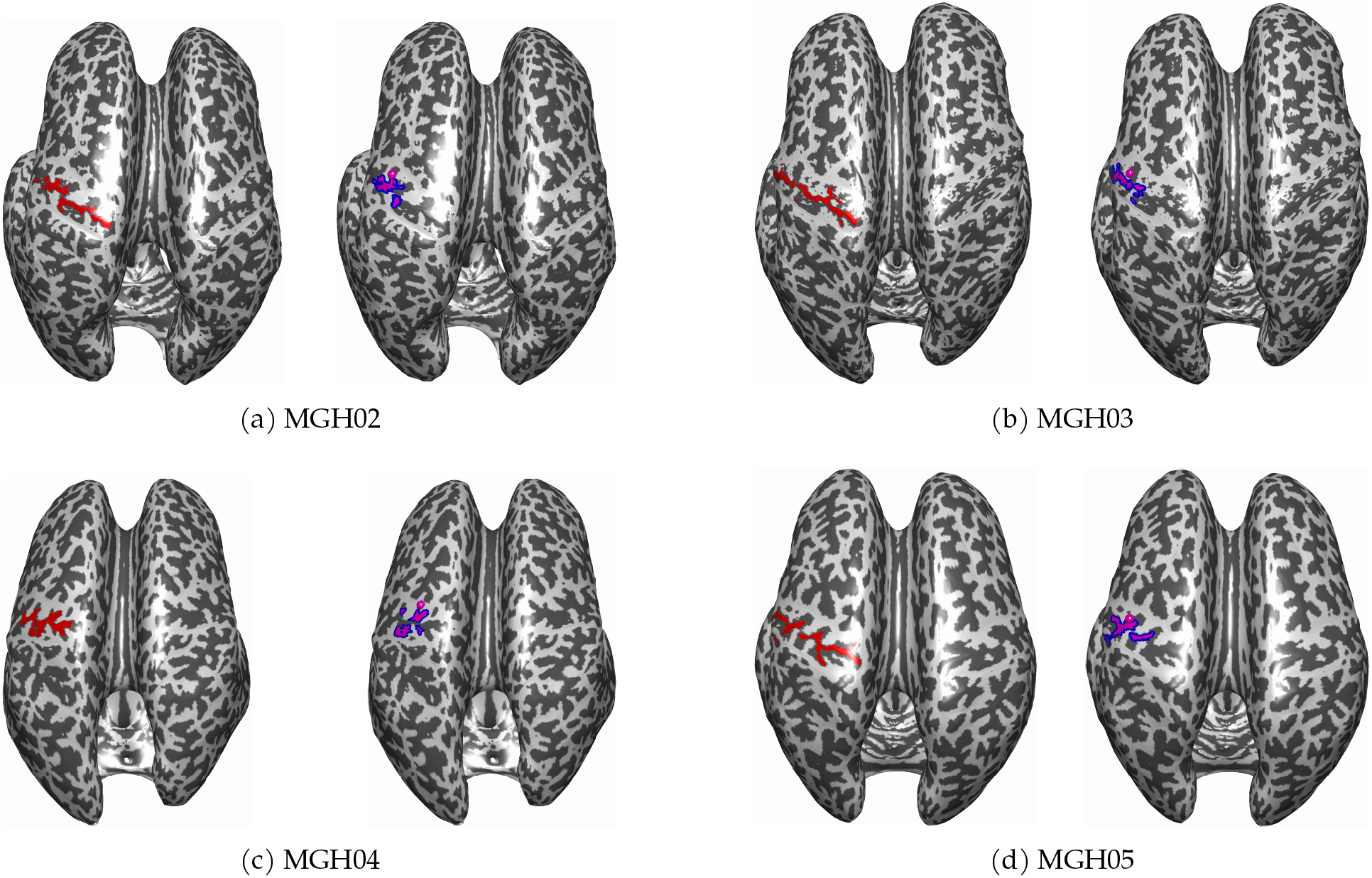
(left) The points on the inflated white matter surface of each subject identifying the central sulcus are shown in red. (right) The tagged point denoting the M1 hand is denoted by the magenta sphere. The target region, shown in magenta, is deterimined by locating the points in the central sulcus nearest to the tagged M1 hand so that 40% of the central sulcus points are covered. The blue region denotes the 2mm distance demarcating the classifier boundary.

**Figure 11.**
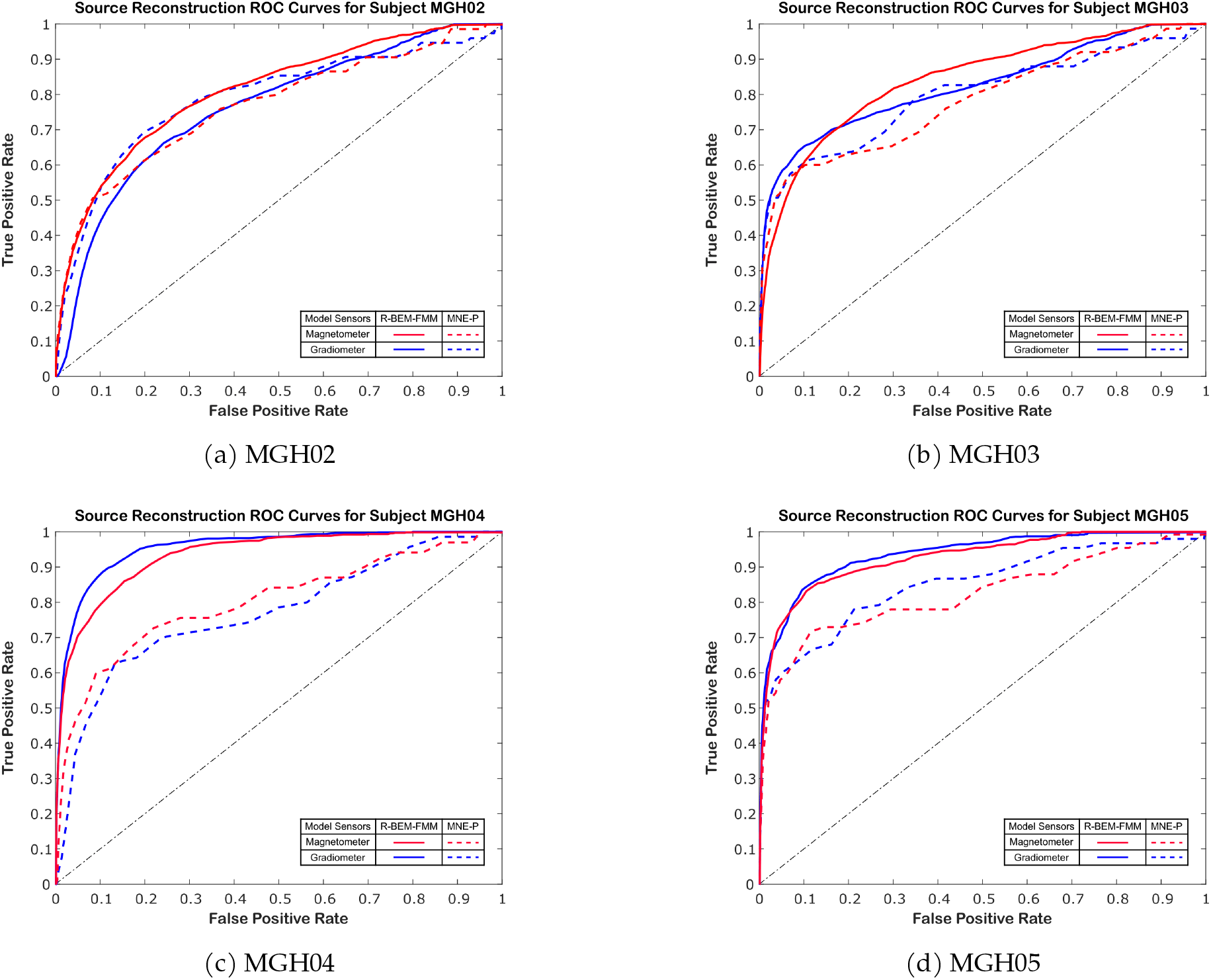
The ROC curves for the source reconstruction results of every subject for both MNE-P (dashed lines) and reciprocal BEM-FMM (solid lines). The red and blue lines denote models which consider only the magnetometer and gradiometer sensor signals, respectively. The black dashed line indicates the “random chance” model.

**Figure 12.**
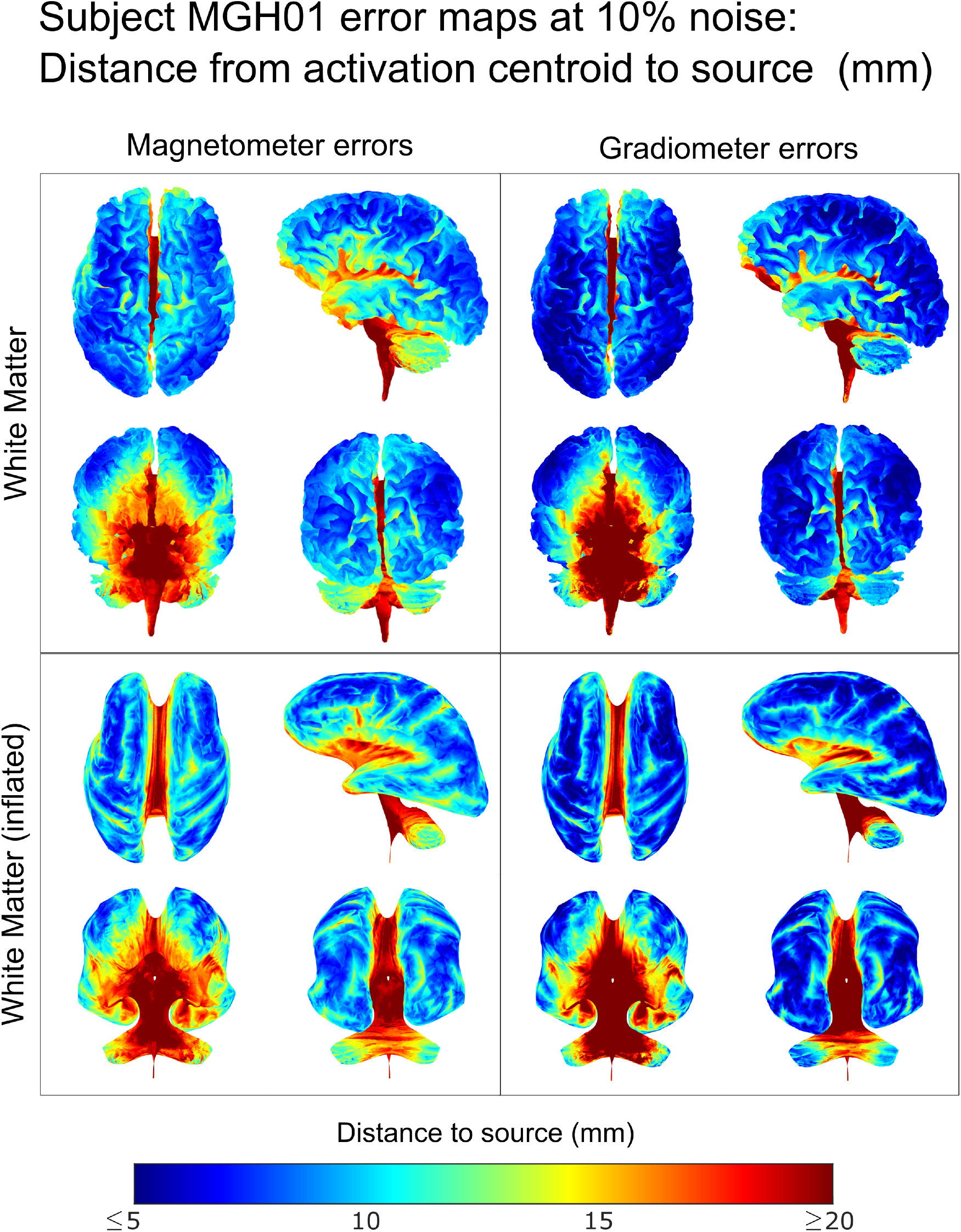
Error maps at 10% noise level (mm) for subject MGH01 on the white matter and inflated white matter surfaces.

**Figure 13.**
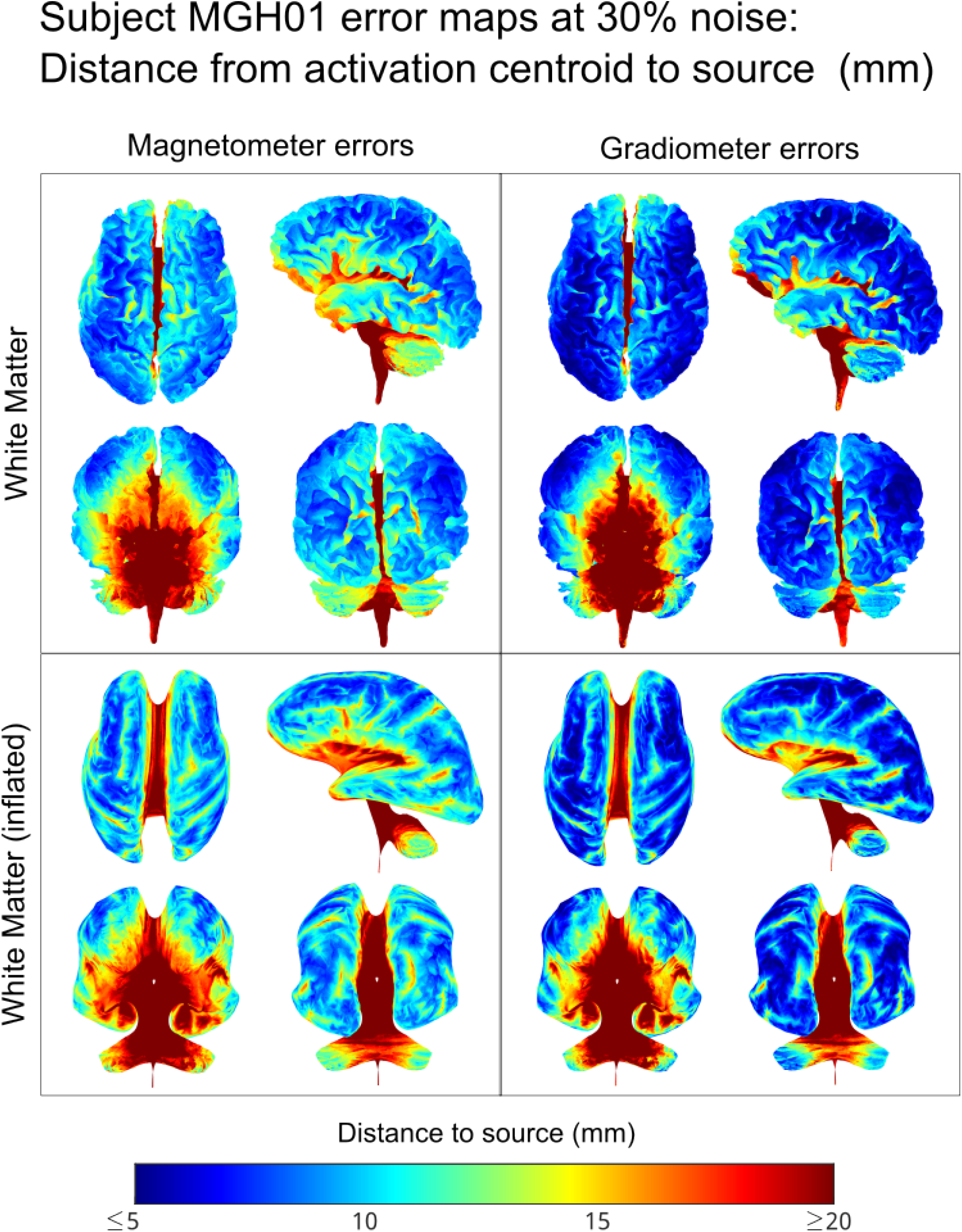
Error maps at 30% noise level (mm) for subject MGH01 on the white matter and inflated white matter surfaces.

**Figure 14.**
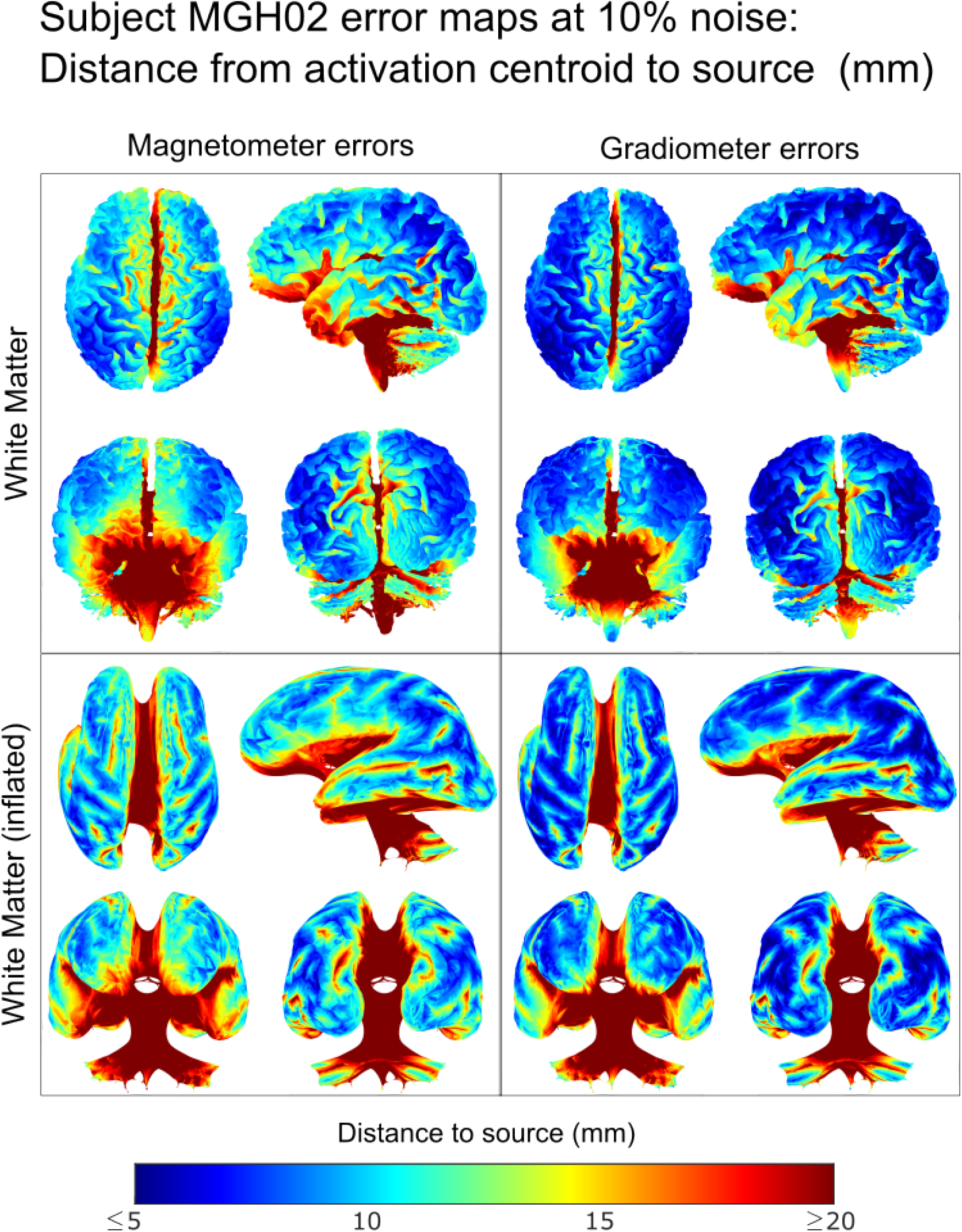
Error maps at 10% noise level (mm) for subject MGH02 on the white matter and inflated white matter surfaces.

**Figure 15.**
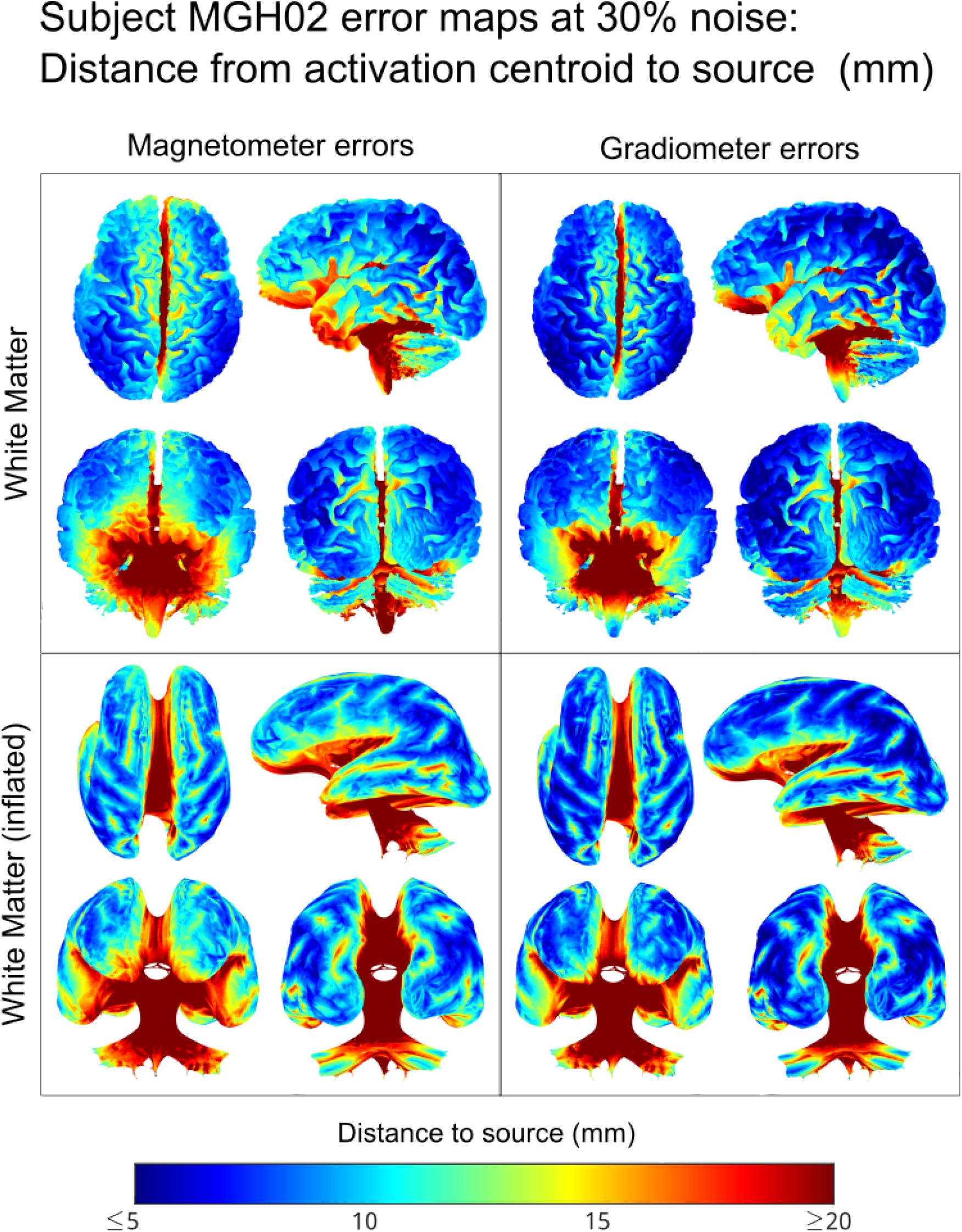
Error maps at 30% noise level (mm) for subject MGH02 on the white matter and inflated white matter surfaces.

**Figure 16.**
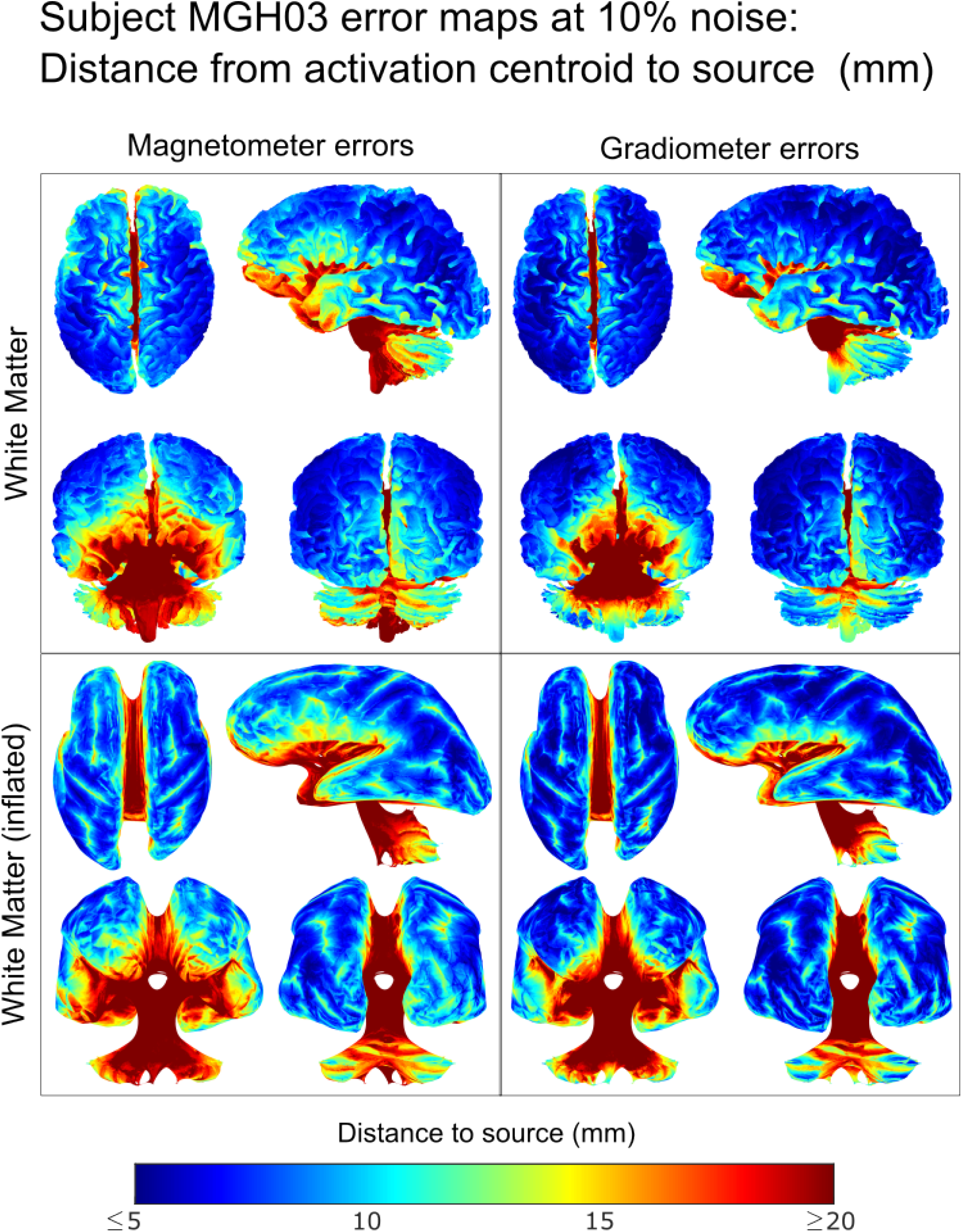
Error maps at 10% noise level (mm) for subject MGH03 on the white matter and inflated white matter surfaces.

**Figure 17.**
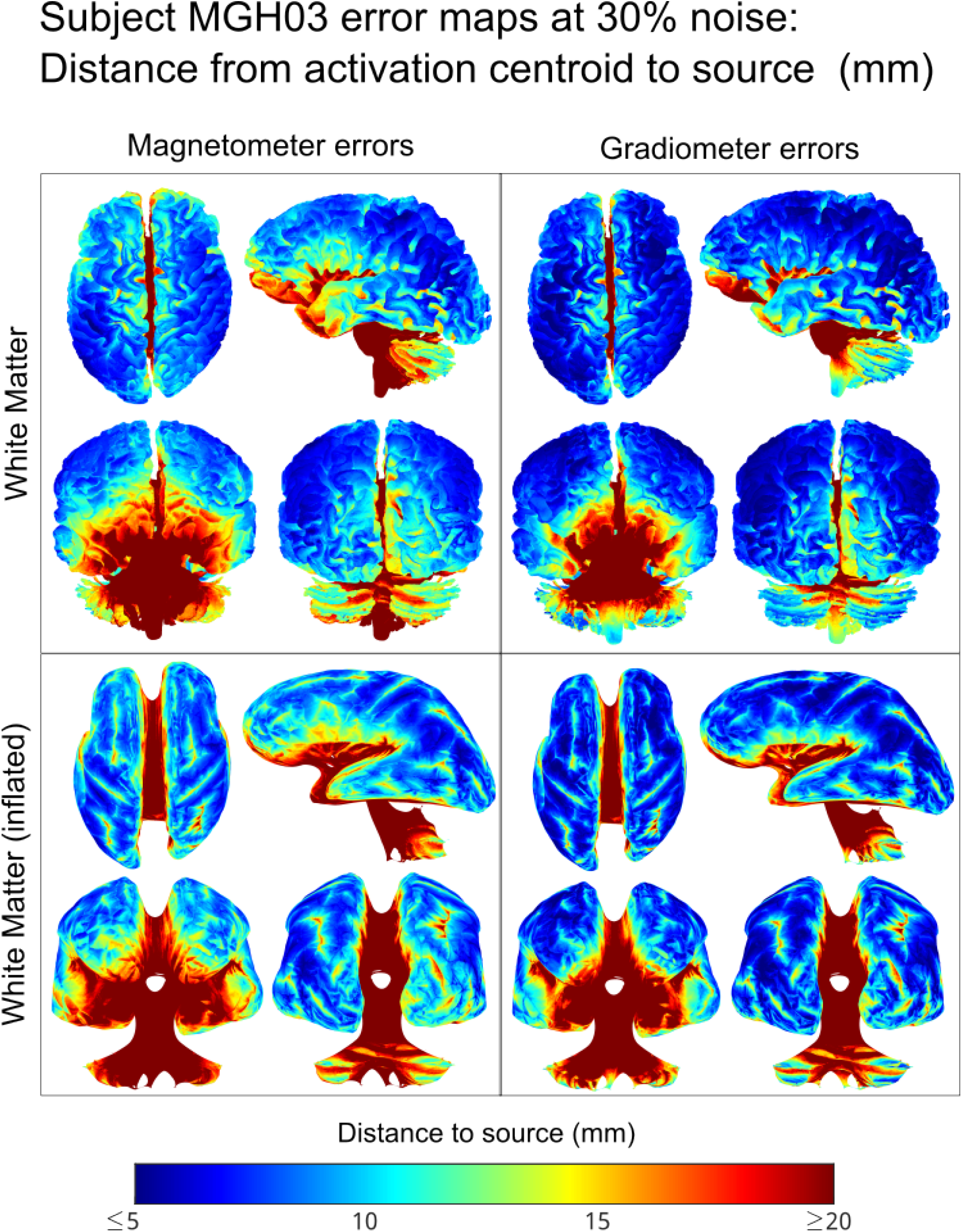
Error maps at 30% noise level (mm) for subject MGH03 on the white matter and inflated white matter surfaces.

**Figure 18.**
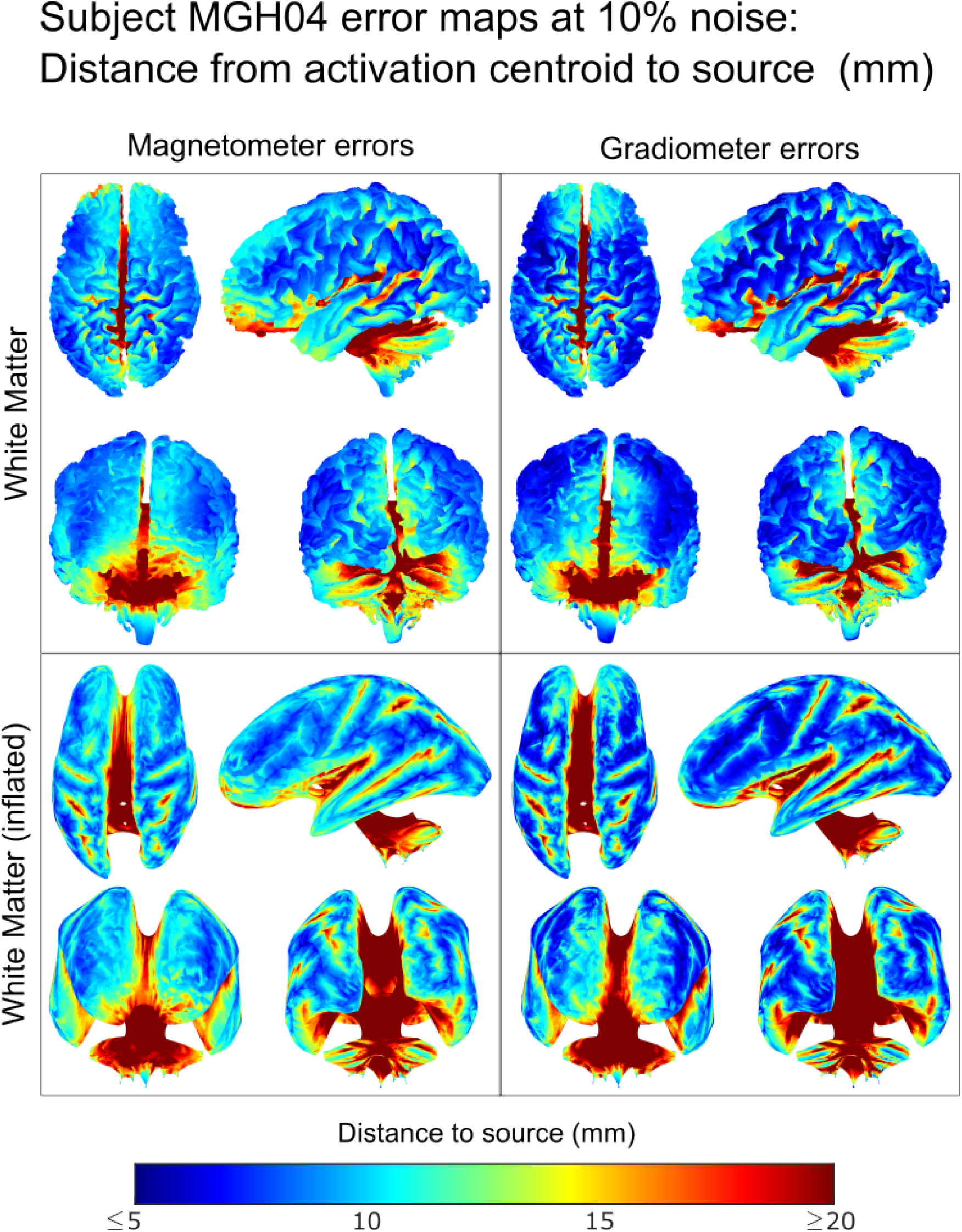
Error maps at 10% noise level (mm) for subject MGH04 on the white matter and inflated white matter surfaces.

**Figure 19.**
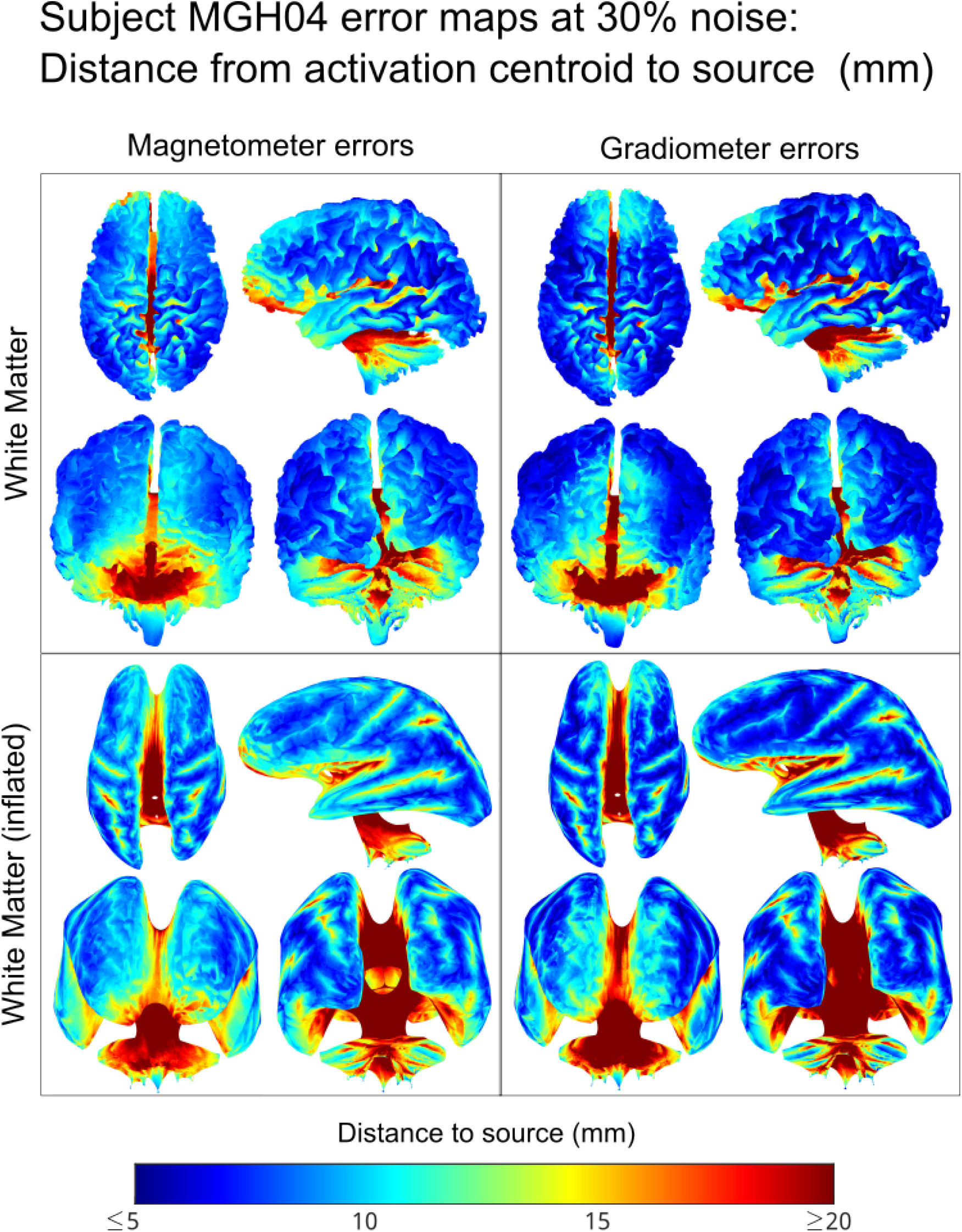
Error maps at 30% noise level (mm) for subject MGH04 on the white matter and inflated white matter surfaces.

**Figure 20.**
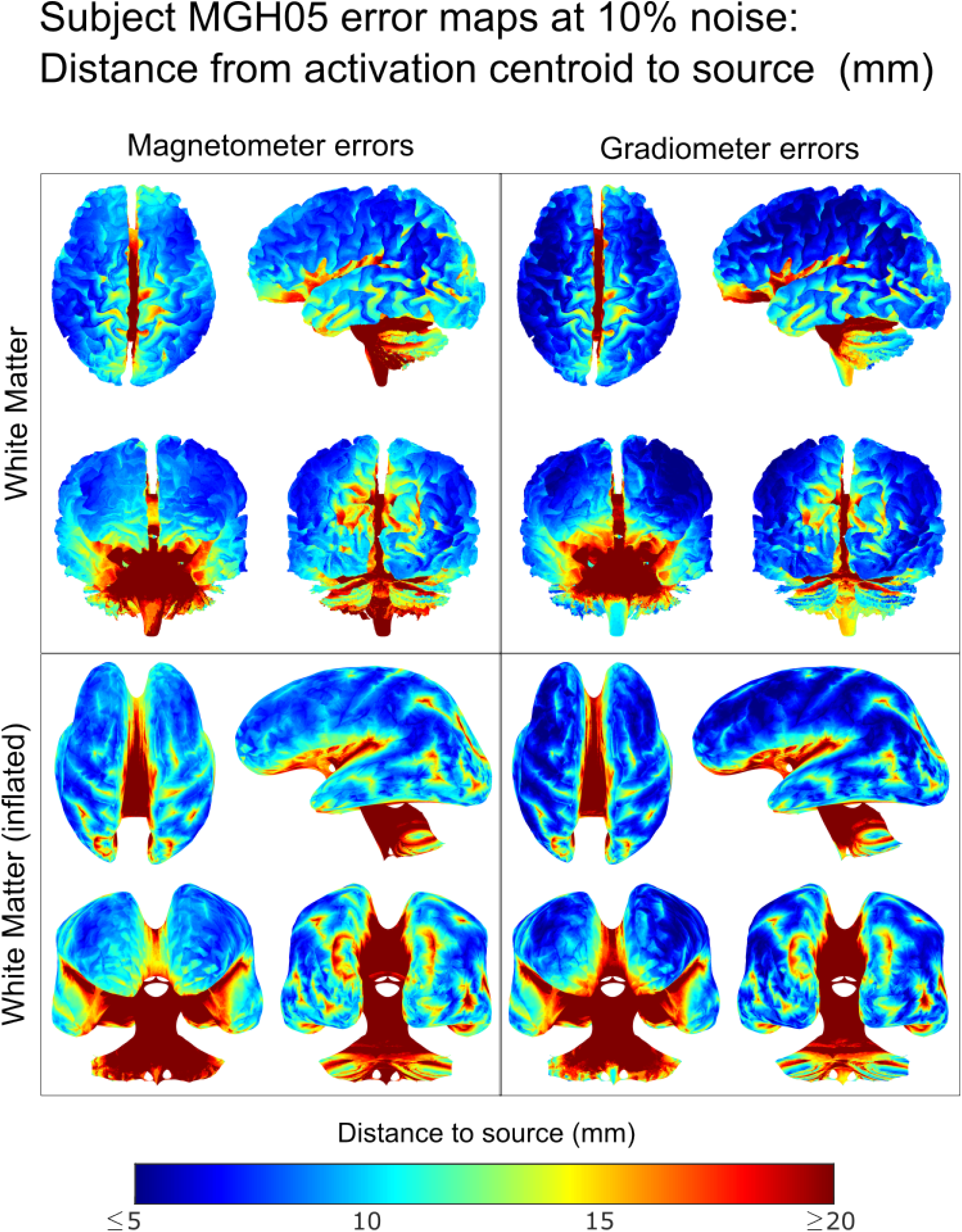
Error maps at 10% noise level (mm) for subject MGH05 on the white matter and inflated white matter surfaces.

**Figure 21.**
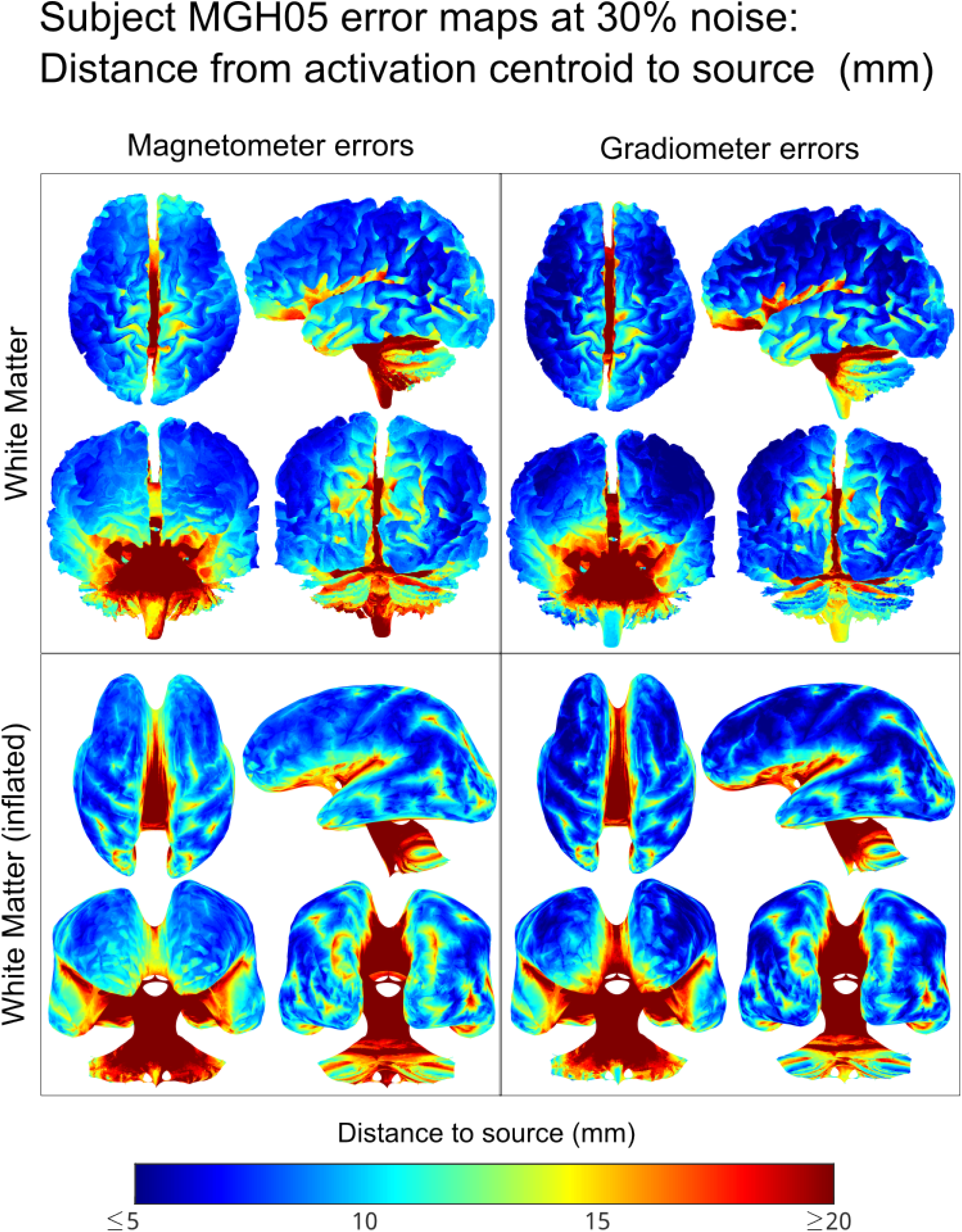
Error maps at 30% noise level (mm) for subject MGH05 on the white matter and inflated white matter surfaces.

## 4. Discussion

### Modeling aspects

Even though we made use of ∼ 250000 sources (WM triangles) in each of our BEM-FMM models, we mention that it is possible to refine and smooth the WM mesh to obtain even denser source spaces. Our decision for the number of sources/triangles we used is motivated by 1) the fact that this the standard number output by `headreco`; and 2) computational constraints imposed by the more extensive direct computations of synthetic data.

We carried preliminary tests with reciprocal solutions using WM meshes of up to 1 million triangles, as well as more recent head models including a higher number of tissue layers (Sim4Life 2025). The computation times for the reciprocal BEM-FMM do not increase prohibitively — further analyses using more detailed models and additional experimental data are interesting directions for future research.

### Analysis of experimental data

From Figures 5 and 6, we see that the reciprocal BEM-FMM source reconstruction at the M20 peak yields activation regions similar to those of MNE-Python. Additionally, we find that activation centroids and peaks are located in similar regions for both reconstruction methods across all subjects. We find that regions of higher activation thresholds appear deep within the Brodmann Area 3b for both source reconstruction methods, consistent with the regions reported in previous experiments (Kawamura et al. 1996; Kakigi et al. 2000).

Because our method makes use of a denser source space, it does not require smoothing and we observe sharper contrasts which follow sulcal wall patterns in the estimated regions of activation. In contrast, MNE-Python requires smoothing, since without it the reconstructed source strength density would appear as a sparse cloud of points. This smoothing process has the effect of blurring areas with isolated peaks. This effect may or may not be desirable, depending on the experiment.

The AUC values in Table 3 indicate good model performance for both reciprocal BEM-FMM and MNE-Python, with our method being slightly favored. However, the results in Table 3 need to be interpreted with care, as we have only an estimated ground truth; and we transformed the problem into a binary classification problem. Additionally, any formal comparison between MNE-Python and our reciprocal BEM-FMM method would require numerical experiments using the same source models and meshes; in this way, the model discrepancies on source estimation of synthetic data can only be attributed to the usage of different forward models. Likewise, the effect of increasing the number of sources (or equivalently, the spatial density of the source space within the cortex) needs to be tested using identical head models. Our results in this paper are meant to be interpreted as a validation for our proposed method. We leave these considerations on the accuracy of inverse methods, with respect to source density, as an interesting direction for future research.

### Evaluation against synthetic data

The results in Table 4 indicate that our inverse method is robust against the addition of increasing levels of noise. Comparing Tables 4a and 4b, one can see that the distance errors associated to 75% threshold activation centroids are, on average, smaller than those associated to source estimation peaks. This effect is also observed in the error maps. It appears that to localize an isolated source, the centroid of the reconstructed region activation may be a more accurate estimate of the true source location than the peak.

The error maps in Figure 9 and Supplement C confirm that source reconstruction of sources in the deepest parts of the cortex can be better recovered from magnetometer data than from gradiometer data. We also observe that sources along the central sulcus are among those with the lowest average errors.

## 5 Conclusions

We proposed, implemented, and tested an accelerated reciprocal solver for MEG source localization based on BEM-FMM. This new method gives an opportunity to employ high-density source spaces in MEG source estimation.. Source localization results for somatosensory evoked fields on 5 healthy subjects indicate activation on the anatomically correct region. Evaluation against synthetic data from dipoles placed on the primary somatosensory cortex shows that our methods perform well under the addition of noise to the source signal, with an average distance error of 6 mm and a standard deviation within a range of 0.03–0.69 mm at different SNR levels.

## Acknowledgments

G.N.P., D.A.D., A.W., G.N., and S.N.M. were supported by the NIBIB Grant 1R01EB035484, and the NIMH Grant 1R01MH130490. T.R. was partially supported by the NINDS grant 1R01NS126337 and

M.H. by NIMH grant 5R01NS104585. J.H. received funding from the German Federal Ministry of Education and Research (BMBF) grant DryPole (01GQ2304A) and the Free State of Thuringia (2018 IZN 004), co-financed by the European Union under the European Regional Development Fund (ERDF).

## Supplements

### A MEG-TMS reciprocity theorem in frequency domain and in time domain

#### A.1 Lorentz reciprocity theorem

The initial reciprocity relation, or Lorentz reciprocity theorem, in integral form without magnetic sources (cf. Equation (A.1) in Heller and Hulsteyn 1992),

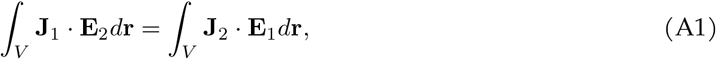

implicitly uses the symmetry of Maxwell’s equations. It is derived from the full set of Maxwell equations, written exclusively in phasor form and without any additional assumptions. By peforming a set of multiplications and subtractions (Plonsey 1972; Balanis 2012), Equation A1 relates vector phasors of two arbitrary sinusoidal impressed current source distributions, **J**_1_ and **J**_2_, operating at the same frequency *ω* to the vector phasors of their respective electric fields, **E**_1_ and **E**_2_. The integration is done over the entire space where the current source distributions exist. According to Lorrain and Corson 1988, the Lorentz reciprocity theorem (e.g. its simplified form in a source-free region) “is paradoxical because it establishes a relation between two unrelated electromagnetic fields”.

#### A.2 Singular impressed harmonic currents in cosine form

In the present application, **J**_1_(**r**) is the impressed phasor point current source located at **r**_1_ within the cortex with vector dipole moment **p**_1_ [A m]. On the other hand, **J**_2_(**r**) is the impressed phasor line current of an MEG sensor (a gradiometer or magnetometer) outside the head operating as a TMS (transcranial magnetic stimulation) coil. Without loss of generality, both independent impressed current sources are assumed to be in phase. This makes it possible to set their phasors as two real numbers simultaneously. The corresponding real-valued impressed current densities can then be written in the form (cf. Equation (A.2) in Heller and Hulsteyn 1992)

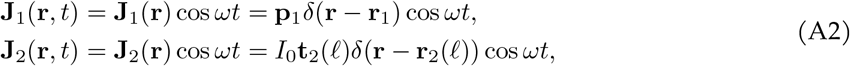

with all constants being real. Here, **r**_2_(ℓ) is a vector running along the contour *C* of the coil conductor (i.e. the MEG sensor) (where ℓ ∈ *C*), **t**_2_(ℓ) is unit tangent to the coil conductor, *I*_0_ is the coil current amplitude [A], and *δ* is the delta function.

#### A.3 Transformation of Lorentz reciprocity equation by expressing E_1_(r)

After substitution of Equation A2 into Equation A1, we obtain (cf. Equation (A.4) in (Heller and Hulsteyn 1992))

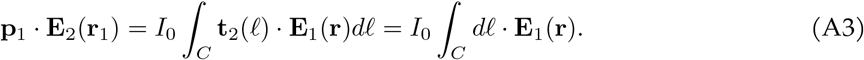

In the quasi-static case, the MEG cortical dipole creates an in-phase conduction current and the associated electric field distribution within the head with the same cos *ωt* dependence, similar to DC steady state. Simultaneously, a time-varying magnetic flux everywhere in space will be generated by this total current distribution in the conducting head. To the highest order of magnitude, the magnetic flux is in phase with the current, again similar to DC steady state. Therefore, we can set for the real-valued magnetic flux **B**_1_(**r**, *t*),

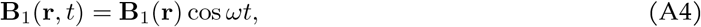

where **B**_1_(**r**) is a purely-real vector phasor. A nonzero circulation of phasor electric field **E**_1_ along the closed contour *L*, or a nontrivial line integral on the right-hand side of Equation A3 is due to Faraday’s laws of induction, which reads in terms of the phasors,

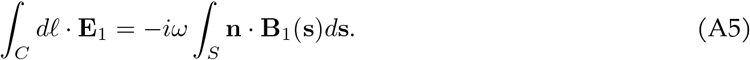

Substitution of Equation A5 into Equation A3 yields (cf. Equations (A.5) and (A.6) of Heller and Hulsteyn 1992),

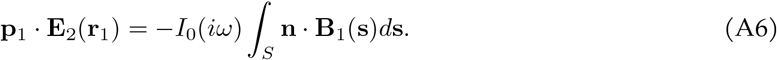

#### A.4 Transformation of Equation A6 by expressing E_2_(r)

In order to satisfy Equation A6 with the real phasor **B**_1_(**r**) and the real constants **p**_1_ and *I*_0_, respectively, the phasor **E**_2_(**r**_1_) of the TMS electric field must be purely imaginary. Let us show that this is really the case.

The TMS coil electric field **E**_2_(**r**) on the left-hand side of Equation A1 and A3 is due to the coil generator current **J**_2_(**r**) in Equation A2. This field has two components: a solenoidal part and a conservative part. In terms of phasor magnetic vector potential **A**_2_(**r**) and phasor electric scalar potential *φ*_2_(**r**), one obtains (cf. p. 261 in Balanis 2012),

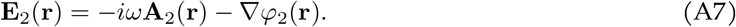

In the quasi-static approximation, the magnetic vector potential is expressed by a static formula through the TMS coil current and without the effect of the conducting head. It is strictly in phase with the coil generator current in Equation A2, so its phasor **A**_2_(**r**) is a real function,

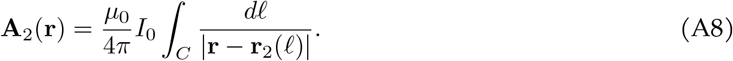

The phasor electric scalar potential *φ*_2_(**r**) is that of electric charges induced at conductivity boundaries in response to the primary solenoidal coil field component, −*iω***A**_2_(**r**). It must therefore be of the same form, −∇*φ*_2_(**r**) = −*iω*∇φ_2_(**r**), with φ_2_(**r**) being real. From Equation A7, one then has,

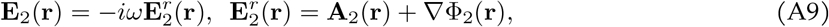

with the real phasor 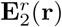.Substituting Equation A9 into Equation A6, one arrives at the final result

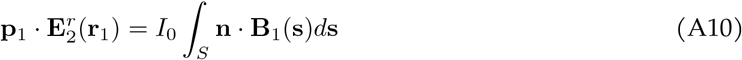

#### A.5 Reciprocity theorem in time domain for harmonic excitation

Multiplying both sides of Equation A10 by cos *ωt*, one can obtain the corresponding time-domain result for harmonic excitation,

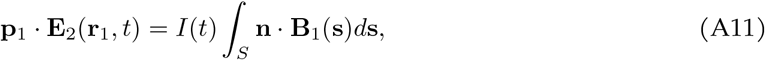

where

– *I*(*t*) = *I*_0_ cos *ωt* is a harmonic TMS coil current; and
– 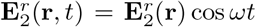 is a harmonic time integral, 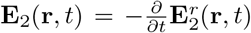,of the TMS coil field **E**_2_(**r**, *t*).

Differentiating Equation A11 over time leads to,

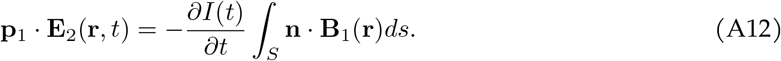

This is a useful result stated previously in (Nummenmaa et al. 2013). It follows in particular 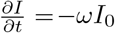 and 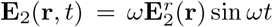.Plugging these into Equation A12 gives us again Equation A11.

### B Derivation of the inverse operator

Here, we give a derivation of the inverse operator *M* of Equation 15 based on error minimization; this is adapted (correcting a minor logical flaw) from the derivation of (Liu 2000).

Let **b** be the vector of measured MEG signals, **x** the vector of dipole strengths, and **n** a noise vector.

Then, the measured MEG signals are related to the dipole strengths **x** by the following equation

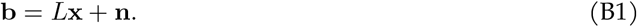

Suppose that the dipole strengths follow a multivariate Gaussian distribution of zero mean and covariance matrix *R* (the source-covariance matrix). Likewise, assume that the noise follows a multivariate Gaussian with zero-mean and covariance matrix *λ*^2^∑ (the noise-covariance matrix, with a regularization parameter). Then, by definition,

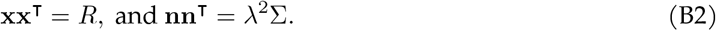

We calculate the linear operator *M* that minimizes the expected error:

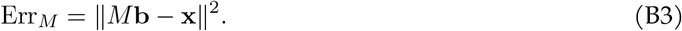

Substitute the expression for **b** of Equation B2 into the expected error to find

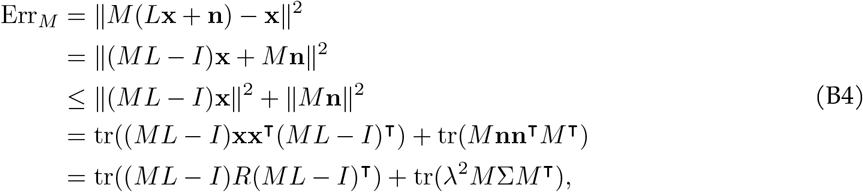

where tr is the trace operator, i.e. the sum of the diagonal elements of the matrix. In particular, we have that the trace is linear and for a matrix *A*, tr(*A*) = tr(*A*^⊺^). Expanding the term ((*ML* −*I*)*R*(*ML* −*I*))^⊺^, we find

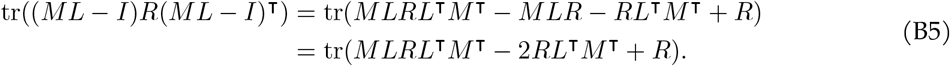

Here, we used the fact that tr(*MLR*) = tr((*MLR*)^⊺^) = tr(*RL*^⊺^*M* ^⊺^). So we have

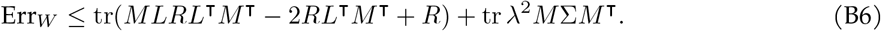

To find the operator *M* that minimizes the error, we take the gradient with respect to *M* in the right-hand-side of the equation above and equate to 0, obtaining:

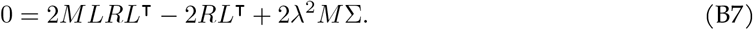

From here, we obtain

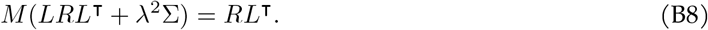

Hence,

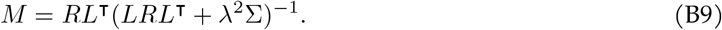

### C Additional Tables and Figures

